# Isolation of Crenothrix bacteria reveals the distinct ecophysiologies of filamentous methanotrophs and adaptations to redox stress

**DOI:** 10.64898/2025.12.18.695142

**Authors:** Kazuhiro Umezawa, Jackson M. Tsuji, Yukinori Tani, Seichi Nohara, Rudolf Amann, Manabu Fukui

## Abstract

At the dawn of modern microbiology, Cohn observed abundant filamentous bacteria in drinking water wells that he named *Crenothrix polyspora*. Subsequent research has revealed the methanotrophic metabolism of Crenothrix bacteria and their disproportionately high activity in stratified lakes compared to unicellular strains, yet laboratory cultivation has proven elusive, leaving the ecophysiology of Crenothrix bacteria largely unknown. Here we report the isolation of two methanotroph strains of the “lacustrine Crenothrix” clade from an iron-rich wetland and reveal their highly unique cell biologies and potential ecological roles. Using physiological approaches, we demonstrate that the strains perform a microaerobic methane metabolism while growing distinct filaments having wide and directionally oriented connective structures. The strains further have broad genomic repertoires for addressing redox stress that we show are uniquely associated with lacustrine Crenothrix compared to related methanotrophs based on genome data. Aligning with laboratory observations, we identify lacustrine Crenothrix bacteria along potential redox gradients in the wetland at iron-rich snow sites, and we further detect such bacteria in diverse global ecosystems based on public metagenome searches. Together, our data strongly point to an ecophysiology of lacustrine Crenothrix bacteria that is tightly linked to redox stress, and we propose these bacteria may uniquely store or share metabolic intermediates via their filamentous lifestyle to thrive under such conditions. Our results provide a fresh view of the diversity, evolution, and ecology of aerobic methanotrophs, connecting over 150 years of microbiology research opening vast new opportunities to probe bacterial adaptations that drive global methane cycling under redox stress.

## Introduction

Methane is a potent greenhouse gas whose emissions are increasing globally from anthropogenic sources and aquatic ecosystems such as wetlands, lakes, and oceans [1, 2]. In freshwater systems, which produce methane equivalent to a quarter of the terrestrial carbon sink [3], aerobic methane-oxidizing bacteria (methanotrophs) that rely on molecular oxygen (O_2_) as a reactant in their metabolism play a crucial role in suppressing methane emissions [4–6]. Yet how these methanotrophic bacteria respond to environmental changes, such as aquatic deoxygenation [7, 8], is unclear, leaving a large knowledge gap for how methane emissions may change or can be managed [5, 9].

Due to the production of methane in anoxic regions, aerobic methanotrophs are commonly found at the boundaries of where O_2_ becomes available in ecosystems. To survive in such O_2_-limited settings and maximize their methane access, methanotrophs require adaptations to manage redox stress. Previous work has reported the use of alternate metabolisms such as denitrification [10] or fermentation [11] by methanotrophs to reduce their O_2_ demands, partnership of methanotrophs with oxygenic phototrophs to utilize cryptic O_2_ [12], and motility of methanotrophs to reach regions with optimal O_2_ levels [13] (reviewed in [9]). Such work has focused on free-living unicellular methanotrophic bacteria despite the occurrence of morphologically diverse forms, such as filamentous methanotrophs, in the environment.

Cohn described filamentous bacteria in drinking water wells over 150 years ago at the dawn of modern microbiology [14]. Although initially suspected to metabolize iron [15], the described species, named *Crenothrix polyspora*, was later confirmed to be methanotrophic using molecular approaches [16]. Subsequent work has detected such filamentous Crenothrix bacteria not only in drinking water and groundwater systems but also in stratified lakes, where they made disproportionately high contributions to methane oxidation compared to unicellular methanotrophs [17]. Other characterized filamentous bacteria employ unique strategies compared to free-living microorganisms to address redox stress, including sheath-based gliding motility to access nutrients [18, 19], storage of metabolic substrates in vacuoles for use under different redox conditions [20, 21], and transport of electrons along filaments to metabolize across redox gradients [22, 23]. Crenothrix bacteria may thus have distinct ecophysiologies and modes for metabolizing methane compared to their unicellular relatives, with important implications for the microbial methane cycle. Yet despite over a century of microbiology research, Crenothrix bacteria have proven elusive to stably cultivate or isolate in the laboratory, leaving the environmental role of these bacteria and their physiologies severely understudied.

In this work, we report the isolation of two methanotrophic bacterial strains from the “lacustrine Crenothrix” clade and provide a detailed description of their cell biology, physiology, ecology, and taxonomy. Our combined cultivation-based, genomic, and field sampling work reveal the unique ecophysiologies of these filamentous methanotrophs, their common association with redox boundary regions, and potentially novel adaptations they may rely on to support their lifestyle compared to unicellular strains. Our work provides important insights into an environmentally relevant form of methanotrophic bacteria and opens crucial new avenues to understand the ecophysiological drivers of the microbial methane cycle.

## Methods

### Isolation of filamentous methanotrophs

Iron-rich microbial mat material was sampled from Oze National Park (Japan) at a wetland site in the park’s western region (site KOJ02; 36°55.025’ N, 139°11.888’ E) in June and August 2022. Following sampling, mat material was inoculated into custom basal cultivation media (see Supplementary Methods and Table S1) in glass serum bottles with a micro-oxic (approximately 5% O_2_) or anoxic headspace. Strains were isolated by serial dilution and sorting of cells using a micromanipulator (Supplementary Methods). Purity was checked by inoculation onto agar plates with 10 times diluted R2A medium.

### Amplicon and genome sequencing

Cells grown in PSVn medium (developed in this study; headspace of 63.3 : 2 :16.7:16.7 dinitrogen : carbon dioxide : methane : air at 1.5 atm pressure) were harvested from turbid cultures by centrifugation. Extraction of DNA from harvested cells was performed using the Wizard Genomic DNA purification Kit (Promega, Madison, WI). Extracted DNA was used to perform amplicon sequencing of the 16S rRNA gene (V4 region) using a MiSeq (Illumina, San Diego, CA) as described in the Supplementary Methods. The same DNA samples were then used for whole genome sequencing. Sequencing libraries for long-read sequencing were prepared using the Native Barcoding Kit 24 V14 (SQK-NBD114.24; Oxford Nanopore Technologies, Didcot, UK), and resulting libraries were each sequenced on three Flongle flow cells (R10.4.1) for 24 h. Short-read sequencing was performed by the Bioengineering Lab Co. (Sagamihara, Japan) using the DNBSEQ-T7 platform (2×150 bp reads; see Supplementary Methods).

Amplicon data was analyzed via QIIME2 version 2024.2 [24, 25], including denoising by DADA2, and hybrid assembly of genome sequencing data was performed using the Hybracter pipeline [26] version 0.7.4 (Supplementary Methods). After assembly, genomes were annotated using DFAST version 1.6.0 [27], and functional genes were predicted using METABOLIC-G version 4.0 [28] and FeGenie version 1.0 [29]. Taxonomic classification was performed using GTDB-Tk version 2.4.0 based on GTDB release 220 [30, 31].

### Physiological analyses

Methane and O_2_ concentrations in culture headspaces were measured over time using a gas chromatograph with a thermal conductivity detector (GC3210S; GL science, Tokyo, Japan) and WG-100 column (GL science). Cultures were grown in PSVn medium under a methane-containing micro-oxic headspace (as above) with 1 mM poorly crystalline ferric oxide particles and 0.1 µM lanthanum(III) nitrate. During analysis, helium was used as the carrier gas (46 mL min^−1^), the oven and injector were heated to 50°C, and the detector was used with an electric current of 120 mA at 55°C.

For transmission and scanning electron microscopy analyses, isolates were grown in PSVn medium as above, with the exception that lanthanum was omitted, until cultures were turbid. Harvested cells were then fixed and prepared for analysis as described in the Supplementary Methods, with sample preparation and observation being performed by Tokai Electron Microscopy, Inc. (Nagoya, Japan). Samples for transmission and scanning electron microscopy were observed under a JEM-1400Plus transmission electron microscope (JEOL, Tokyo, Japan) and JSM-7500F scanning electron microscope (JEOL), respectively.

We additionally performed time-lapse observations of the strain AF98 culture during growth. The culture was inoculated in PSVn medium (without ferric oxide particles but with 0.1 µM lanthanum(III) nitrate) on a glass Petri dish, and the petri dish was placed into a custom gas-tight glass chamber. A gas mixture (50 : 25 : 25 of dinitrogen : methane : air) was continuously provided at 10 mL min^−1^ to the chamber headspace, and the chamber was incubated at 20-25°C under an inverted microscope (IX71; Evident, Tokyo, Japan). Phase-contrast images of growing cells were taken every 30 seconds.

### Comparative genomics and phylogenetics

Genome sequences of selected *Methylococcaceae* members were obtained from the National Center for Biotechnology Information (NCBI) database together with the genome of an outgroup *Methylococcales* member. The resulting genome set, including strains AF98 and MI19235, contained 38 genomes (or 37 excluding the outgroup strain). A phylogenomic tree was then built using GToTree version 1.8.3 [32] based on 170 concatenated marker proteins as described in the Supplementary Methods. A 16S rRNA gene phylogeny was constructed to complement this analysis. Sequences were aligned using MAFFT version 6.864, and a maximum likelihood phylogenetic tree was built, based on 1260 conserved nucleotide sites, using the T3p+G+I evolutionary model with 100 bootstrap iterations via MEGA11 [33–35]. Selected environmental 16S rRNA gene sequences from public databases, including those classified as *Crenothrix* in the Silva database (release 138.2) [36], were included in an extended phylogenetic analysis. The sequences were aligned using MAFFT and used to construct a maximum likelihood phylogeny via MEGA12 [37] with 1189 conserved nucleotide sites. The TN93+G+I evolutionary model was used with 1000 bootstrap iterations.

The 38 genomes were annotated using METABOLIC-G version 4.0 [28] and FeGenie version 1.0 [29]. In addition, whole protein-coding genes in the genomes were classified into homologous groups using OrthoFinder version 2.5.5 [38] for further gene identification. Potential bacteriohemerythrins and hemerythrin domain-containing proteins were identified based on a conserved protein domain family for hemerythrin (ID cl15774) using the Batch Web version of the CD-search tool [39]. Methanol dehydrogenases XoxF and MxaF were identified by phylogenetic placement. Sequences were aligned via MAFFT with the reference sequence set used by Wu and colleagues [40], and aligned sequences were used to construct a maximum likelihood phylogenetic tree, based on 500 conserved amino acid sites, using MEGA11 with 100 bootstrap iterations and the LG+G evolutionary model. To identify genes encoding nitrite reductases (*nirS* and *nirK*), the reference dataset used by Pold and colleagues [41] was searched via hmmsearch version 3.4 using an e-value threshold of 10^−100^ [42].

Orthologous gene groups were further compared between genomes via an ordination. Counts of orthologous groups per genome were normalized as proportions within each genome. Singleton orthologous groups (i.e., with only one gene member across all genomes) were excluded from calculations. The resulting proportion table was used to calculate a dissimilarity matrix via the diversity.beta_diversity function in scikit-bio version 0.6.2 [43] using Bray-Curtis distances. The stats.ordination.pcoa function in scikit-bio was then used to generate the ordination by principal coordinate analysis (PCoA). Groupings of genomes by clade were tested for statistical significance using a permutational analysis of variance (PERMANOVA). The stats.distance.permanova function of scikit-bio was used with 10,000 permutations.

To complement the above analyses, expanded sets of 99 and 313 genomes associated with the *Methylococcales* order were also downloaded and analyzed to perform a pan-core analysis and semi-automated searches of denitrification genes and hemerythrin domain-containing genes. Analysis is described in the Supplementary Methods.

### Collection of environmental samples

To understand the distribution of filamentous methanotrophs in the environment and their potential activity, we surveyed the wetland site across summer, fall, and spring of 2022 and 2024. In summer and fall, we targeted sites with robust iron mats, and we compared these to sites having iron material mixed with plant matter (“iron residue” sites) and sites outside the iron-rich zone of the park (“non-iron” sites). In spring, we sampled snow overlying iron-rich sites and took either snow cores or surficial snow samples where snow was in the final stages of spring melt. Samples for DNA extraction were collected as bulk material or by filtration and were preserved with DNA/RNA Shield (Zymo Research; California, U.S.A.; details in Supplementary Methods). Samples were then kept cool until return to the laboratory, where they were frozen. For CARD-FISH analyses of iron-rich snow cores, material was collected in sterile bags, allowed to melt, and transported cool to the laboratory for fixation. Collection of physicochemical parameters, including total dissolved iron concentrations, is described in the Supplementary Methods.

### Amplicon sequencing of environmental samples

Extraction of DNA was performed using the collected environmental material or filter membranes using the ZymoBIOMICS DNA Mini Kit (D4300; Zymo). Amplicon sequencing of the 16S rRNA gene (V3-V4 region) was then performed for the samples by the Bioengineering Lab Co. (Sagamihara, Japan; see Supplementary Methods). QIIME2 version 2024.10 [24] was used to analyze the amplicon sequencing data, including primer trimming, sequence denoising, and taxonomic classification based on the Silva r138 SSU database [36]. Sequences were further classified into potential *Crenothrix* clades by phylogenetic placement as described in the Supplementary Methods.

An ordination was constructed for the sequenced environmental samples. Samples were rarefied to the lowest sequence count value in the dataset (i.e., 26,686) using the feature-table rarefy module of QIIME2. Following this, scikit-bio version 0.6.2 [43] was used to construct the ordination as a PCoA distance biplot as described in the Supplementary Methods. A PERMANOVA was performed, to test if grouping samples by wetland site type was statistically robust, using the same methods as for comparative genomics data above.

Relative abundances of *Methylococcales* members were compared between iron mat samples and other wetland samples using scipy version 1.16.1 [44] in Python version 3.12.4. A single representative sample from each site and sampling season was selected for the test (see Supplementary Data 2). Given that the assumption of normality was violated based on the scipy stats.normaltest function, a Mann-Whiteney *U* test was used for the comparison via the stats.mannwhitneyu function.

### Methane and carbon dioxide flux measurements

Concentration changes of methane and carbon dioxide were measured using a GasScouter (G4301; PICARRO, Santa Clara, CA) with a static chamber on site. For terrestrial or shallow wetland sites, the chamber was placed on the surface of the soil/sediment or sedge/moss material, whereas the chamber was placed on the water surface for pond sites. Measurements were performed multiple times on each site. For each measurement, the molar fraction of methane and carbon dioxide was corrected based on effects of water vapor, and rates of concentration change were estimated over at least 60 s measurement time. Gas flux was estimated from the resulting rate.

### CARD-FISH analysis

Collected snow core samples were fixed in 4% paraformaldehyde in phosphate-buffered saline (PBS; 130 mM NaCl, 10 mM sodium phosphate, pH=7.3) overnight at 4°C. Fixed samples were then washed twice in PBS and stored in a 1:1 mixture of PBS:ethanol at −20°C. Probes sequences and formamide concentrations, including for the Mγ669 probe used to target *Methylococcales* members, are described in the Supplementary Methods. An aliquot of fixed sample, mixed in a 1:1 ratio with 0.1% agarose, was spread on a gelatin-coated slide glass prepared according to [45]. Subsequent CARD-FISH preparation was performed according to [46, 47]. Cell walls were permeabilized by incubating the spread samples in 10 mg ml^−1^ lysozyme (dissolved in 0.1 M Tris-HCl, 0.05 M EDTA, pH=8.0) for 35 min at 37°C. Endogenous peroxidase was inactivated by incubation in a solution of 0.1 M HCl at room temperature for 1 min, followed by washing with PBS for 1 min, incubation with 3% H_2_O_2_ for 10 min, and washing with PBS and water for 5 min and 1 min, respectively. CARD signal amplification was performed at 46°C using Alexa488-labelled and Alexa594-labelled tyramide. Visualization of samples was performed using a LSM780 laser scanning microscope (Zeiss, Jena, Germany) equipped with an Airyscan detector.

### Global sequence survey

The Sandpiper database, version 1.0.1 [48], was queried for metagenomes containing members of the “*Ca*. Allocrenothrix” (i.e., UBA4132) genus, the *Crenothrix* genus, and the *Methylococcales* order. Only metagenomes having with at least 0.01% relative abundances of “*Ca*. Allocrenothrix” or *Crenothrix* members were retained. In addition, metagenomes that seemed to be associated with laboratory cultures or synthetic microbial communities (based on “taxon”) were filtered from the analysis. The “taxon” entries for metagenomes were then summarized and collapsed into taxon groups to aid visualization (Supplementary Data 3).

## Results and Discussion

### Isolation of filamentous methanotrophs

Sampling iron-rich mat material from the wetland at the sharp redox boundary between the oxic surface and anoxic underlying conditions allowed us to cultivate novel methanotrophs. Incubating the collected material under reduced O_2_ or anoxic conditions in a custom medium with methane and ferric iron, we observed growth of long filamentous bacteria that commonly associated with iron particles. Ferric iron could eventually be removed from the medium in later subcultures, and cultures that were started under anoxic conditions could be grown under 5% O_2_ with higher growth rates. After enrichment by filtration, micromanipulation, serial dilution, and multiple transfers to a refined selective medium (see Supplementary Methods), we obtained two independent axenic cultures of filamentous methanotrophic bacteria, named strains AF98 and MI19235. We validated the purity of the cultures via 16S rRNA gene amplicon sequencing (detection limit: 0.004%) and inoculation in dilute R2A medium.

### Methane-oxidizing physiology and cell biology

Previous attempts to establish cultures of filamentous methanotrophs have resulted in cultures with weak growth that could not be sustained beyond a few subcultures [49]. By contrast, strains AF98 and MI19235 exhibit robust growth as axenic cultures (Fig. 1, Fig. S1), with both strains having a 1-2 day doubling time and maximum observed culture densities of approximately 10^7^ cells mL^−1^. Axenic cultures have remained stable for more than one year of repeated subcultivation. When grown under 5% O_2_ with methane, strains uptake methane until O_2_ is depleted, with a O_2_:methane uptake ratio of approximately 1.0-1.4 that is well below the theoretical 2:1 stoichiometry of O_2_:methane in dissimilatory aerobic methane oxidation (Fig. 1A-B). The low O_2_:methane uptake ratio implies these strains supplement their methanotrophic metabolism with alternative metabolic pathways, such as fermentation [11, 50], to conserve O_2_. The robust growth of these filamentous strains under methane and low O_2_ concentrations facilitated study of their physiology and cell biology in the laboratory.

**Fig. 1.**
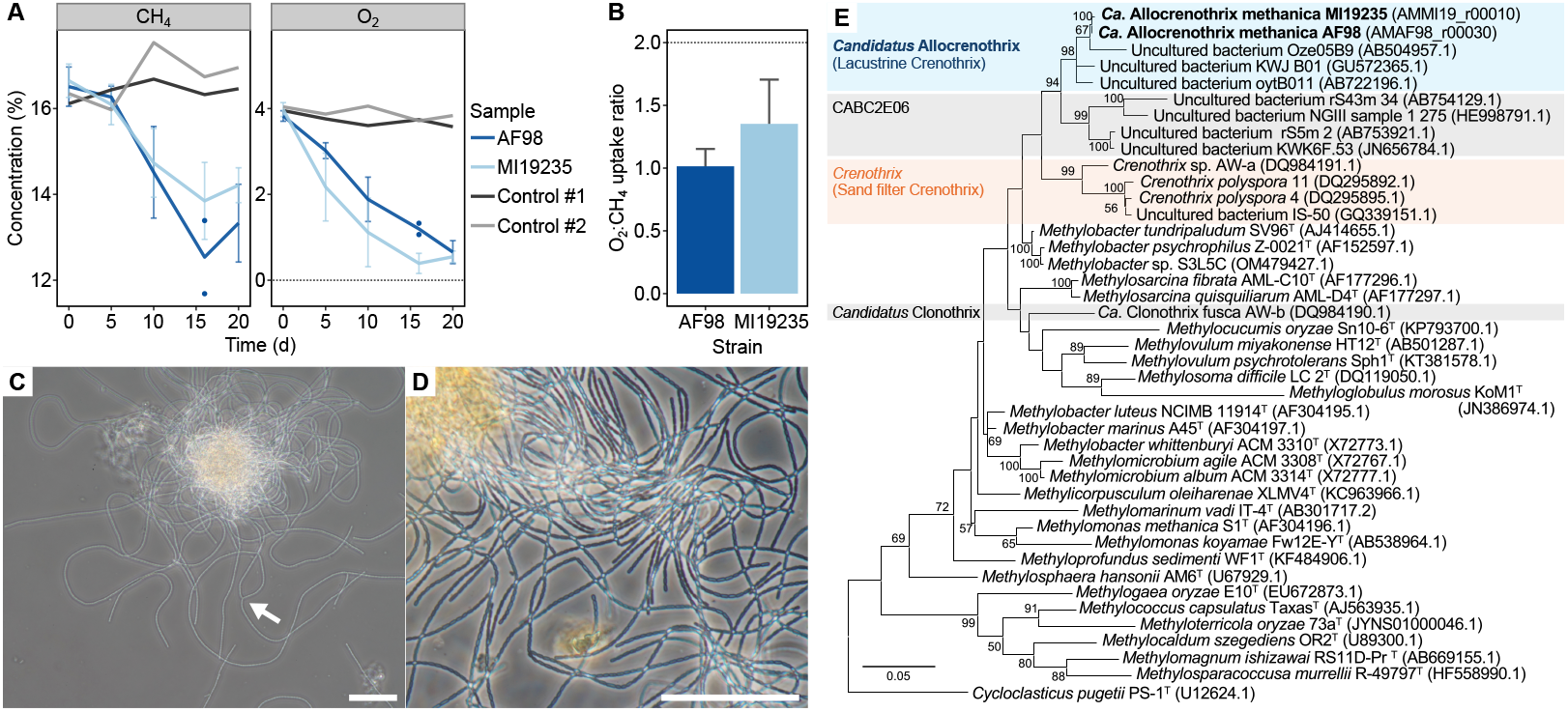
Cultivation of filamentous methanotrophs. (A) Oxidation of methane by strains AF98 and MI19235 compared to abiotic controls. Standard deviations of the mean are shown for biological replicates as error bars (n=3). (B) O_2_:methane uptake ratios for cultures in (A) until complete O_2_ depletion (20 and 16 days for AF98 and MI19235, respectively; error bars are as in A). (C,D) Phase contrast microscopy images of strain AF98 grown under micro-oxic conditions with methane. The arrow (C) shows an example branching structure. The culture in (D) additionally included ferric iron particles. Scale bars represent 50 µm. (E) Maximum likelihood 16S rRNA gene phylogeny of *Methylococcaceae* family members. The phylogeny was based on 1,260 nucleotide sites with 100 bootstrap iterations. Bootstrap values over 50/100 are shown.

Most filaments of strains AF98 and MI19235 are linear and extend in length over the course of incubation, with filaments composed of rod-shaped cells of 1.0-1.3 x 1.9-4.3 µm for strain AF98 (Fig. 1C-D) and slightly larger cells of 1.1-1.5 x 2.5-5.0 µm for strain MI19235 (Fig. S1A). Although filaments of strain MI19235 are generally fragmented, filaments of strain AF98 can extend to hundreds of micrometers in length and occasionally develop branched structures, which we observe more frequently in fully grown cultures (Fig. 1C,; Supplementary Videos 1-3). Tips of filaments are prone to adhesive attachment to surfaces or other filaments (e.g., Supplementary Video 3, bottom left), and we commonly observe tight bundles of filaments for strain AF98 during growth (Fig. 1C). *Crenothrix polyspora* has been observed to produce spore-like structures called gonidia [14, 51], but we do not observe such structures for strains AF98 and MI19235. We also have not observed gliding motility for either strain, although we did observe a gradual serpentine motion of filaments that appeared distinct from Brownian motion of surrounding particles (Supplementary Videos 1-3). We confirmed based on 16S rRNA gene sequences that strains AF98 and MI19235 are members of the gammaproteobacterial *Methylococcales* order, which includes Crenothrix bacteria and other aerobic methanotrophs (Fig. 1E; see below).

At the sub-cellular level (Fig. 2) inside filaments (Fig. 2A, Fig. S1B), we observed folded intracytoplasmic membrane stacks that are typical for aerobic methanotrophs via transmission electron microscopy (Fig. 2B-C, Fig. S1C). We did not observe gas vacuoles, which may be used for buoyancy by some free-living methanotrophs [9]. Between cells in filaments, we identified unusual conjugation structures, where cell membranes apparently open directly into adjacent cells (Fig. 2D-E, Fig. S1C). These conjugations always occurred in a single directional orientation in filaments and were accompanied by a veil-like structure enrobing the connection (Fig. 2B,D,E). Less frequently, we also observed such conjugations on the long sides of cells, potentially resulting in cell pairs or branching filaments (Fig. S1D-E).

**Fig. 2.**
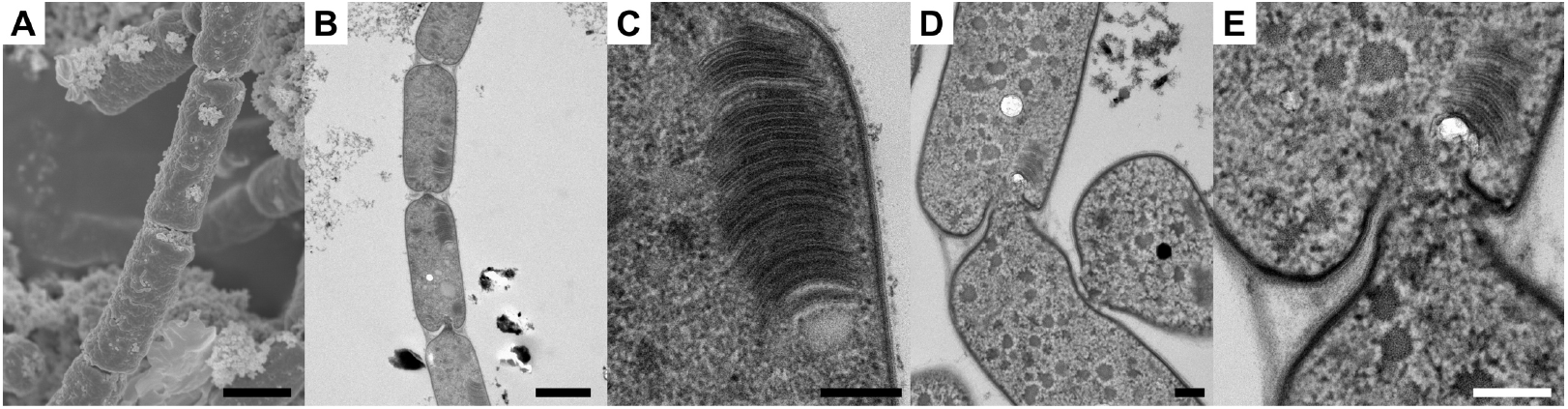
Sub-cellular physiology of methanotroph filaments. Images of strain AF98 are shown when grown under micro-oxic conditions with methane and ferric iron particles (see Fig. S1 for images of strain MI19235). (A) Scanning electron microscopy image of a representative cellular filament. (B,C) Transmission electron microscopy images of connected cells in a filament (B) and their internal structures (C; image taken from the center area of B). (D,E) Transmission electron microscopy image of a cell-to-cell conjugation (E is taken from the center area of D). Scale bars represent 1 µm (A-B) and 0.2 µm (C-D).

Cytoplasmic bridges were consistently over 100 nm in diameter (Fig. 2E), implying they could facilitate movement of cytoplasmic contents between cells. Together, our observations clearly establish strains AF98 and MI19235 as filamentous aerobic methanotrophs capable of growth under low O_2_, with a lack of gonidia and presence of unique cellular structures implying these strains represent a distinct form of Crenothrix bacteria compared to those reported previously.

### Phylogenetic placement

We sequenced and annotated closed circular genomes of strains AF98 and MI19235 to understand their evolutionary placement (Fig. 3). The two genomes with total lengths of 4,134,542 bp and 4,298,823 bp for AF98 and MI19235, respectively, consist of a single circular chromosome and several circular plasmids (8 for AF98 and 5 for MI19235). Each strain has three identical 16S rRNA gene copies per chromosome (Supplementary Data 1). Known Crenothrix bacteria can be subdivided into multiple polyphyletic clades within the *Methylococcaceae* family of the *Methylococcales* order [52], including the “sand filter Crenothrix” clade containing *Crenothrix polyspora* and a “lacustrine Crenothrix” clade containing filamentous methanotrophs observed in stratified lakes [17]. Previous studies have also reported a third clade represented by the filamentous methanotroph *Clonothrix fusca*, which was available in a poorly growing enrichment culture but has not been reported on in nearly 20 years [49, 53]. Based on a 16S rRNA gene phylogeny (Fig. 1E, Fig. S2) and concatenated protein phylogeny (Fig. 3A), strains AF98 and MI19235 fall into the lacustrine Crenothrix clade alongside the uncultured *Crenothrix* sp. D3 [17]. Classification into the lacustrine Crenothrix clade is supported by classification according to the Genome Taxonomy Database (GTDB; release 220), which places strains AF98 and MI19235 in the uncultured UBA4132 genus represented by strain D3 [30].

**Fig. 3.**
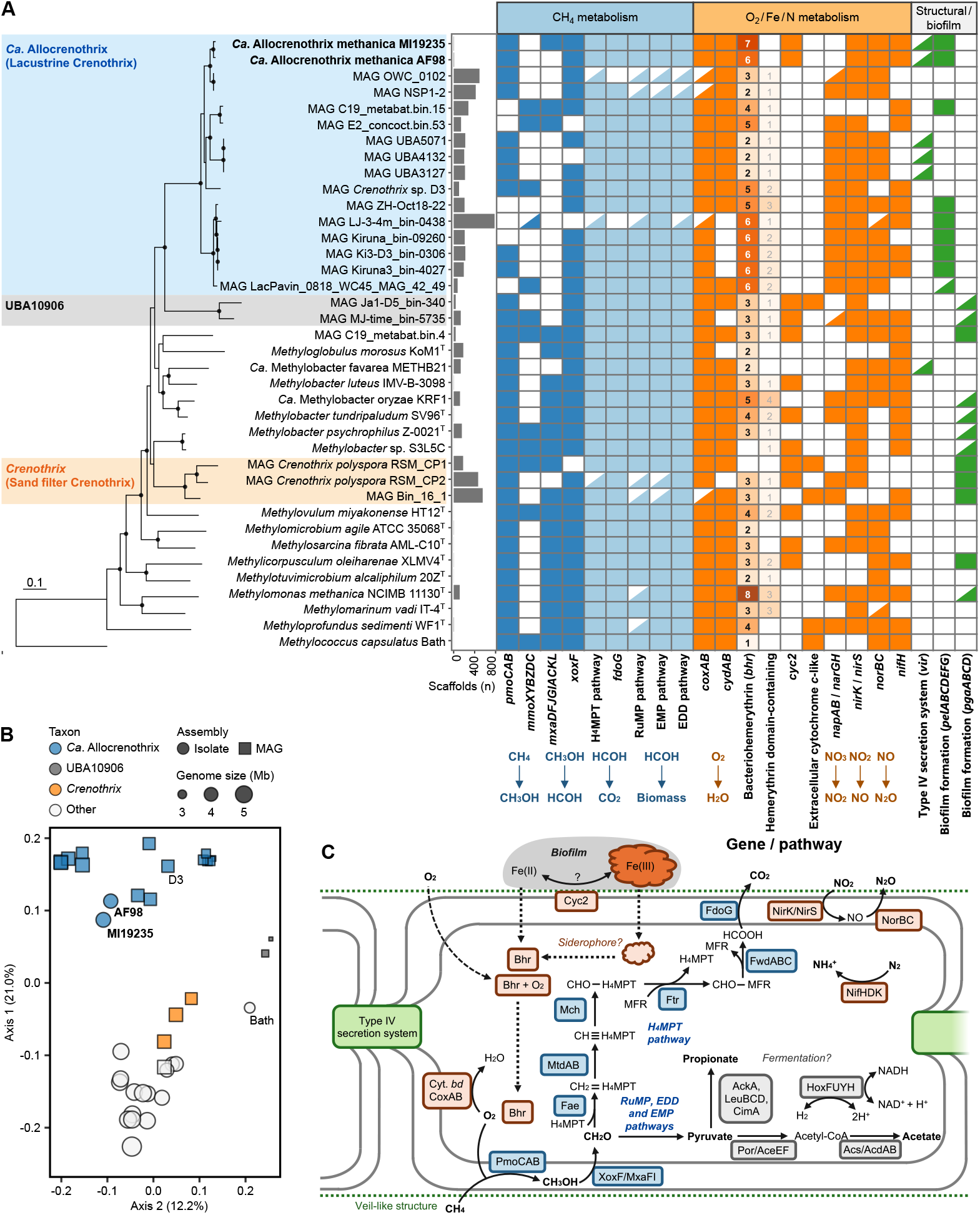
Genomic characterization of “*Ca*. Allocrenothrix” members. (A) Comparative genomic analysis of members of the *Methylococcaceae* family. The maximum likelihood phylogeny (left) was constructed based on 170 concatenated marker proteins with 1000 ultrafast bootstraps. Bootstrap values over 90/100 are shown as black dots. In the gene heatmap (right), solid squares indicate presence of a gene/pathway, solid triangles indicate presence of over half the genes in a pathway, and open squares indicate absence of a gene or presence of less than half the genes in a pathway. Numbers on solid squares for (bacterio)hemerythrin-related genes, along with the intensity of shading, indicates gene copy number. Additional genome statistics and locus information for detected genes are shown in Supplementary Data 1. (B) Ordination of genomes in (A) based on counts of orthologous gene groups in each genome. (C) Cell diagram showing the proposed metabolism of strains AF98 and MI19235.

Strains AF98 and MI19235 share an average nucleotide identity of 95.3% and a 16S rRNA gene sequence identity of 99.8%. This, together with their related physiologies, leads us to propose them as two distinct phylotypes of the same novel lacustrine Crenothrix species [54]. For reasons of priority, we maintain the name *Crenothrix* for the genus described for Cohn’s *Crenothrix polyspora* and propose to name AF98 and MI19235 as strains of “*Candidatus* Allocrenothrix methanica” (Allocrenothrix, i.e., “other Crenothrix”; see etymology below), with UBA4132 (i.e., lacustrine Crenothrix) proposed as “*Candidatus* Allocrenothrix”. We retain *Candidatus* status for the species because the proposed type strain AF98 is being prepared for deposition into culture collections. Although the physiology of classical *Crenothrix* (i.e., sand filter Crenothrix) and *Clonothrix* species have been studied to some degree in the laboratory [16, 49, 51], the lacustrine clade (i.e., “*Ca*. Allocrenothrix”) has been little characterized outside of environmental genomics and microscopy, which further highlights the novelty of strains AF98 and MI19235.

### Comparative genomics

We analyzed the functional gene contents of “*Ca*. Allocrenothrix methanica” strains AF98 and MI19235 to compare them to other cultured and uncultured potential methanotrophs. Our analyses focused on members of the *Methylococcaceae* family given the inclusion of all known Crenothrix bacteria in this lineage. We found that the AF98 and MI19235 genomes, along with most metagenome-assembled genomes (MAGs) classified to “*Ca*. Allocrenothrix” from public databases, encode *pmoCAB* genes for a particulate methane monooxygenase (pMMO) along with genes for the classical pathway of aerobic methane oxidation (Fig. 3A and Note S1; all genes are listed by locus in Supplementary Data 1). At the same time, in place of the *mxaF* gene for calcium-dependent methanol dehydrogenase, many “*Ca*. Allocrenothrix” members encode the *xoxF* gene for the lanthanide-dependent enzyme [55] (Fig. S3), and we speculate that dependence on lanthanides could be one reason for previous challenges in cultivating these filamentous methanotrophs.

“*Ca*. Allocrenothrix” members commonly have ample genomic potential for addressing redox stress. Along with genomic potential for aerobic respiration, including the *coxAB* genes encoding a *caa*3-type cytochrome *c* [56, 57], the majority of “*Ca*. Allocrenothrix” members (including AF98 and MI19235) encode at least five homologs of bacteriohemerythrin, a non-heme-based O_2_ binding protein implicated in O_2_ storage in gammaproteobacterial methanotrophs [58–62] (Fig. 3A, Table S2). By contrast, the genomes we analyzed of cultured unicellular *Methylococcaceae* members generally encode three or fewer of these genes. These analyses were robust when extended to a representative set of over 300 genomes/MAGs from the *Methylococcales* order, with “*Ca*. Allocrenothrix” members having a median hemerythrin domain-containing protein count of 8 compared to a median of 1-4 in other analyzed *Methylococcales* lineages (Table S2). The majority of analyzed “*Ca*. Allocrenothrix” members, including strains AF98 and MI19235, also encode genes for partial denitrification of nitrite to nitrous oxide, including a *nirK* or *nirS* homolog for nitrite reductase and *norBC* homologs for nitric oxide reductase [63] (Fig. 3A). Some “*Ca*. Allocrenothrix” members (not AF98/MI19235) additionally encode *narGHI* homologs for nitrate reductase. Although denitrification genes are commonly encoded by isolated unicellular *Methylococcaceae* members, the gene pathway is absent in all MAGs of classical *Crenothrix* members (Fig. 3A, Table S2). The genomes of strains AF98 and MI19235 also encode potential pathways for fermentation of pyruvate to acetate and/or propionate (Note S1), which could potentially facilitate the low O_2_:CH_4_ uptake ratios of these strains. In addition, homologs of *cyc2*, which could play a role in extracellular electron transfer, were encoded by both genomes, although the exact role of *cyc2* remains unclear [64], and the gene was absent in other “*Ca*. Allocrenothrix” members (Note S1).

To explore other genes potentially unique to filamentous methanotrophs, we performed a pan-core analysis of “*Ca*. Allocrenothrix”, its sister lineage UBA10906, *Crenothrix*, and other *Methylococcaceae* members (Fig. S4). When separated in an ordination based on counts of orthologous gene groups in genomes, members of “*Ca*. Allocrenothrix” formed a separate cluster from other clades (PERMANOVA, n=38, *F*=6.18, *p*=0.001), which further supports that the “*Ca*. Allocrenothrix” lineage is genomically distinct from its sibling lineages (Fig. 3B, Fig. S4A). Aside from core genes common to *Methylococcaceae* members, less than 4% of orthologous gene groups were shared between “*Ca*. Allocrenothrix” and *Crenothrix* members, and none of these genes had clear roles related to cell division or cell structure based on annotation data (Fig. S4B-C; genes are summarized in Supplementary Data 1). Given their phylogenetic separation and unique functional gene profiles, it is possible that *Crenothrix* and “*Ca*. Allocrenothrix” were shaped by distinct evolutionary pressures and could have even evolved filamentous morphology independently.

Among orthologous gene groups that were unique to “*Ca*. Allocrenothrix” members, we detected a *pelABCDEFG* gene cluster, associated with biosynthesis of extracellular polysaccharides involved in biofilm formation [65–69], that was conserved in strain AF98, MI19235, and several other members of the genus (Fig. 3A). This was distinct from the *pgaABCD* genes implicated in biofilm formation that we detected in *Crenothrix* members and other methanotrophs (Fig. 3A). Beyond this, we detected a cluster of *vir* genes, encoding a Type IV secretion system, that were encoded by several “*Ca*. Allocrenothrix” members including strains AF98 and MI19235 (Fig. 3A, Note S1). We speculate that production of extracellular polysaccharides may be important for scaffolding and attachment of filamentous cells, for example allowing cells to remain at locations with optimal concentrations of O_2_ and methane, and Type IV secretion systems could potentially play a role in the cell conjugations we observed (e.g., Fig. 3C) [70]. Together, our analyses establish “*Ca*. Allocrenothrix” as a genomically distinct lineage of aerobic methanotrophs with enhanced genomic potential for coping with redox stress compared to related clades.

### Detection in iron-rich wetlands

To clarify the potential ecologies of “*Ca*. Allocrenothrix” members, we surveyed the subalpine wetlands from which strains AF98 and MI19235 were isolated and performed amplicon sequencing, microscopy-based analyses, and gas flux measurements (Fig. 4). Along with non-iron rich peatlands, the wetlands contain abundant iron-rich zones that can be subdivided into extreme iron-rich mats from which both strains were cultured, iron-rich mat residue mixed with plant material, and iron-rich snow material that forms over iron mat regions in winter (Fig. S5A-C). Iron-rich snow regions can be temporally distinguished as pre-melt material and snow in the final stages of spring melt, where a reddish precipitate appears on the snow surface in a phenomenon locally called “Akashibo” [71]. Iron-rich mats and their overlying snow were commonly stained an intense reddish-brown colour and could develop high dissolved iron concentrations, which reached up to 0.3 mM in surficial mat samples, highlighting the potentially steep redox gradients between oxic and anoxic wetland zones in these regions (Supplementary Data 2; see also [71, 72]).

**Fig. 4.**
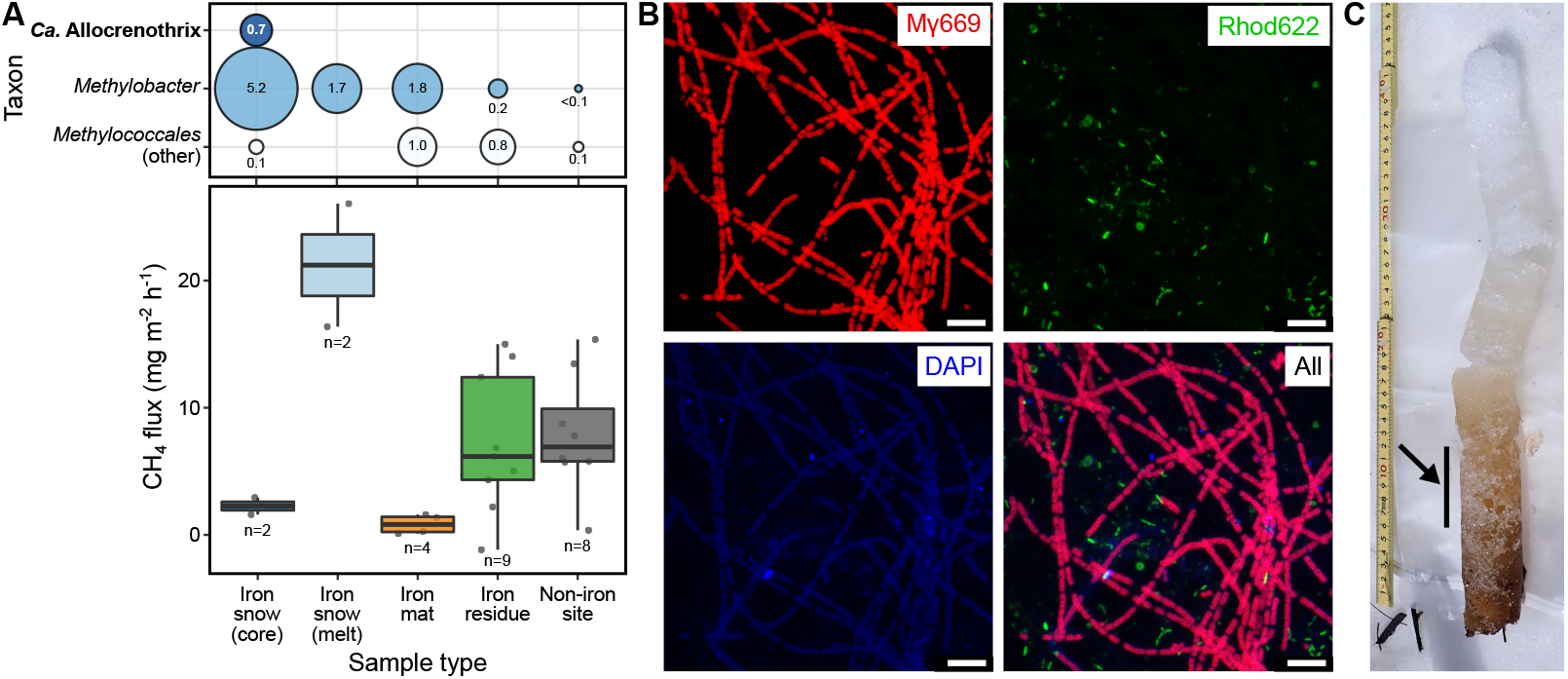
Ecology of filamentous methanotrophs in iron-rich wetlands. (A) Gas fluxes and potential methanotroph communities from wetland samples. The bubble plot (top) shows the maximum relative abundances of *Methylococcales* clades detected from each wetland site type based on 16S rRNA gene amplicon data. The boxplots (bottom) show the methane fluxes measured from distinct wetland sites and seasonal timepoints within each site type. Amplicon and flux data per site/timepoint are shown in Fig. S6, and raw data are available in Supplementary Data 2. (B) Microscopy image of iron-rich snow core material (from site KOJ20, 6-12 cm above surface; see below) using the CARD-FISH assay. Results are shown for the Type I methanotroph probe Mγ669 (red), DAPI staining (blue), and the Rhod622 probe targeting *Rhodoferax* members as a control (green). Scale bars represent 10 µm. Additional CARD-FISH images are shown in Fig. S8. (C) Iron-rich snow core from the KOJ20 site. The arrow indicates the snow core region used for CARD-FISH analyses shown in (B).

Collecting samples across multiple seasons and years, we observed large differences in methane fluxes (Fig. 4A and Fig. S6). Iron-rich snow cores (2.9 and 1.6 mg methane m^−2^ h^−1^, n=2) and iron mat zones (0.83 mg methane m^−2^ h^−1^ ± 0.76 S.D., n=4) had consistently low measured methane fluxes despite overlying anoxic sediments, while iron residue (7.21 mg methane m^−2^ h^−1^ ± 5.52 S.D., n=9), and non-iron rich sites (7.91 mg methane m^−2^ h^−1^ ± 4.73 S.D., n=8) had higher but variable methane fluxes. Meanwhile, zones in the final stages of snow melt had the highest observed methane fluxes (16.4 and 26.1 mg methane m^−2^ h^−1^, n=2), potentially due to rapid methane release during spring thaw or disruption of microbial communities. Differences between wetland site types were evident at the microbial community level, with an ordination of 16S rRNA gene amplicon data (Fig. S7) indicating that iron-rich snow, iron mat, iron residue, and non-iron rich samples formed statistically robust groupings (PERMANOVA, n=25, *F*=3.32, *p*=0.0001). Along with lower methane emissions, iron-rich snow core and iron mat samples also had elevated relative abundances of *Methylococcales* members (3.06% ± 2.05 S.D., n=6), based on amplicon data, compared to samples from other sites (0.24% ± 0.48 S.D., n=16; Mann-Whitney *U*=93.0, *p*=0.00057, two-sided). Together, these gas flux and amplicon data suggest that iron-rich snow and iron mat zones could be key sites for aerobic methanotrophy in the wetland.

Within the detected *Methylococcales* sequences, we identified “*Ca*. Allocrenothrix” members based on manual reclassification of amplicon data by phylogenetic placement (Fig. 4A; see Supplementary Methods). Although we did not detect “*Ca*. Allocrenothrix” members in iron-rich mat samples from which strains AF98 and MI19235 were isolated (detection limit: 0.007%), we identified multiple “*Ca*. Allocrenothrix” sequence variants from snow core samples, including one sequence variant that exactly matched the sequences of strains AF98 and MI19235 and had relative abundances of up to 0.6% (sequence oze-d3cbca2; see Supplementary Data 2). Sequences of “*Ca*. Allocrenothrix” members were also identified from an iron-rich snow sample that was collected from the wetland over 15 years ago (GenBank AB504957.1; Fig. 1E) [71], which suggests that “*Ca*. Allocrenothrix” members have long-term environmental relevance within the methanotroph communities in this region. We always detected other *Methylococcales* members in samples with “*Ca*. Allocrenothrix” sequences, and “*Ca*. Allocrenothrix” members made up approximately 8-15% of the potential gammaproteobacterial methanotroph community (Fig. S6). By contrast, we did not detect classical *Crenothrix* members in any analyzed sample.

We further performed microscopic single cell identification and quantification of iron-rich snow core material using fluorescence in situ hybridization (FISH; Fig. 4B-C). By staining with the *Methylococcales* probe Mγ669, whose sequence exactly matches that of strains AF98 and MI19235 except for a single base on the 3’ end, we observed long Mγ669-stained filaments that likely represent “*Ca*. Allocrenothrix” members along with free-living Mγ669-stained cells (Fig. 4B-C, Fig. S8A-B). The filamentous arrangements of the visualized cells were analogous to TEM observations of strains AF98 and MI19235. Samples were heterogeneous, and the Mγ669-stained cells varied in their morphology, yet filamentous methanotrophs consistently appeared to be important *Methylococcales* members in these iron-rich snow samples. Ferrous iron concentrations of up to 0.45 mM have been measured from interstitial water nearly 30 cm above surface within snow cores from the wetland [71], implying these iron snow regions can exist at the interface between oxic and reducing conditions while receiving fluxes of methane from underlying sediments. That strains AF98 and MI19235 exactly match the dominant “*Ca*. Allocrenothrix” sequence variant from these systems suggests that adaptations observed in these strains are representative of those employed by filamentous methanotrophs in the environment.

### Global ecology

We surveyed publicly available metagenome data to understand the broader distribution of “*Ca*. Allocrenothrix” members (Fig. 5). Searches for classical *Crenothrix* members were used as a baseline for our analyses. Querying over 700,000 metagenome samples via the Sandpiper database [48], we uncovered 1427 samples that included “*Ca*. Allocrenothrix” (i.e., UBA4132) members, compared to 156 samples for *Crenothrix* members (Fig. 5A).

**Fig. 5.**
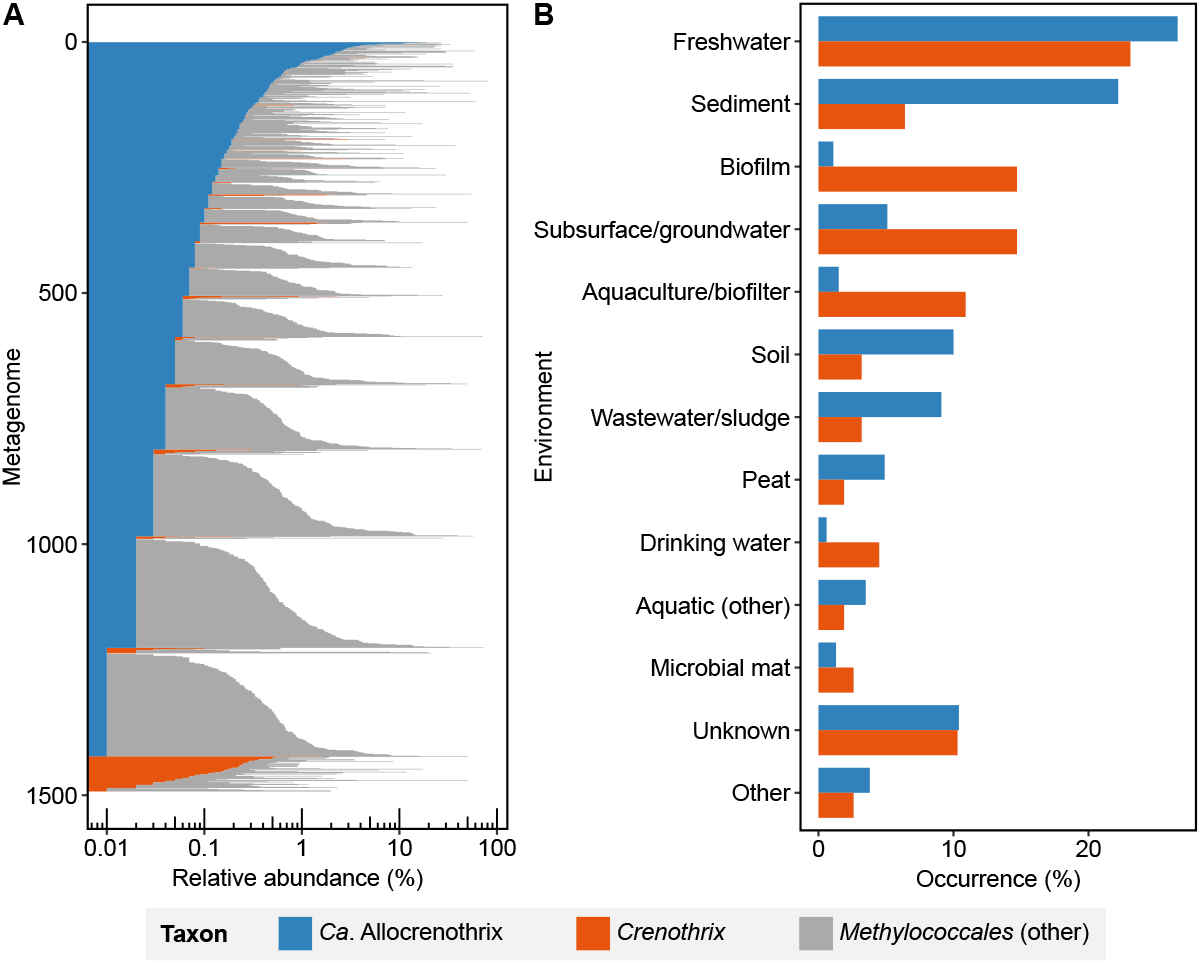
Global distribution of potential filamentous methanotrophs. (A) Relative abundances of “*Ca*. Allocrenothrix” and classical *Crenothrix* members in publicly available metagenome samples. Samples containing at least 0.01% of members of these genera are shown. (B) Environmental affiliations (by “taxon”) of metagenomes where “*Ca*. Allocrenothrix” and *Crenothrix* members were detected. Taxa are collapsed into higher-level environmental groups (see Supplementary Data 3), and groups are shown if they included at least 1.5% of metagenomes where a methanotroph taxon was detected. Other groups are collapsed as “Other”.

Comparing the environmental (“taxon”) attributions of metagenomes from which these genera were identified, we found that members of both genera were commonly detected in freshwater samples, but “*Ca*. Allocrenothrix” members were more frequently associated with sediment, soil, wastewater/sludge, and peat samples than *Crenothrix* members (Fig. 5B; Supplementary Data 3). By contrast, *Crenothrix* members were more frequently associated with biofilm, subsurface/groundwater, aquaculture/biofilm, and drinking water samples (Fig. 5B). Despite known accuracy limitations of “taxon” attributions of metagenomes, results for *Crenothrix* members match literature reports of these bacteria in oligotrophic environments such as drinking water and groundwater [14, 15], implying these data capture real trends. Widespread detection of “*Ca*. Allocrenothrix” members in environments that typically include redox gradients, such as sediments and sludges, is consistent with genomic and field survey data that indicate these organisms commonly encounter redox stress.

### Ecophysiology linked to redox stress

Despite their first observation over 150 years ago, filamentous methanotrophs have remained severely understudied in their ecological roles due to a lack of available laboratory cultures. Our isolation of two filamentous methanotroph strains and analysis of their physiology and complete genome sequences provide fundamental new insights into ecophysiology of this elusive bacterial group. Given that “*Ca*. Allocrenothrix methanica” strains AF98 and MI19235 grow at reduced O_2_ with low O_2_:methane uptake ratios, that these strains encode high numbers of redox stress-linked genes compared to methanotrophs in related clades, and that close relatives of these strains are detected in methane-containing and potentially reducing environments, our data strongly indicate that “*Ca*. Allocrenothrix methanica” and other species in the “*Ca*. Allocrenothrix” genus are highly adapted to life under redox stress. These results match previous detection of “*Ca*. Allocrenothrix” (i.e., lacustrine Crenothrix) members in stratified lake systems [17]. Moreover, our results indicate that large genomic differences exist between filamentous methanotroph groups, with classical *Crenothrix* (i.e., sand filter Crenothrix) members potentially having distinct evolutionary adaptations and ecological preferences from the “*Ca*. Allocrenothrix” group.

Although the mechanisms driving the redox stress-linked ecophysiology of “*Ca*. Allocrenothrix” remain to be explored in detail, our cell biology observations and genomic findings allow us to propose potential adaptations that may allow these filamentous methanotrophs to inhabit low O_2_ environments (Fig. 6). Although strains AF98 and MI19235 do not possess gliding motility, they are prone to surface attachment and have relatively large cytoplasmic volumes compared to unicellular methanotrophs, as observed for “*Ca*. Allocrenothrix” members in the environment [17]. Rather than relying on motility like certain unicellular methanotrophs, we thus propose that “*Ca*. Allocrenothrix” members could anchor via biofilms at regions with optimal methane and O_2_ concentrations, and cells could then use stored compounds, such as O_2_-bound bacteriohemerythrins, to persist under periods of transient redox change such as O_2_ depletion (Fig. 6A). Strains AF98 and MI19235 also possess unique cellular connective structures, and it is natural to speculate that these structures could play a role in movements of compounds between cells, even if by diffusion. Given that “*Ca*. Allocrenothrix” members can be detected near redox gradient regions, we speculate that sharing of metabolic intermediates, such as methanol produced during the O_2_-dependent step of aerobic methane oxidation [73], between cells in filaments could promote their activity by supplying O_2_-deficient cells with carbon and energy sources (Fig. 6B). These proposed models to “store” or “share” metabolic intermediates could potentially enhance both persistence and activity of “*Ca*. Allocrenothrix” cells in O_2_-limited niches and represent novel redox adaptation strategies compared to those described previously for unicellular methanotrophs. With cultures of strain AF98 and MI19235 now available for study, these models and their implications for global methane cycling can now be investigated in laboratory research.

**Fig. 6.**
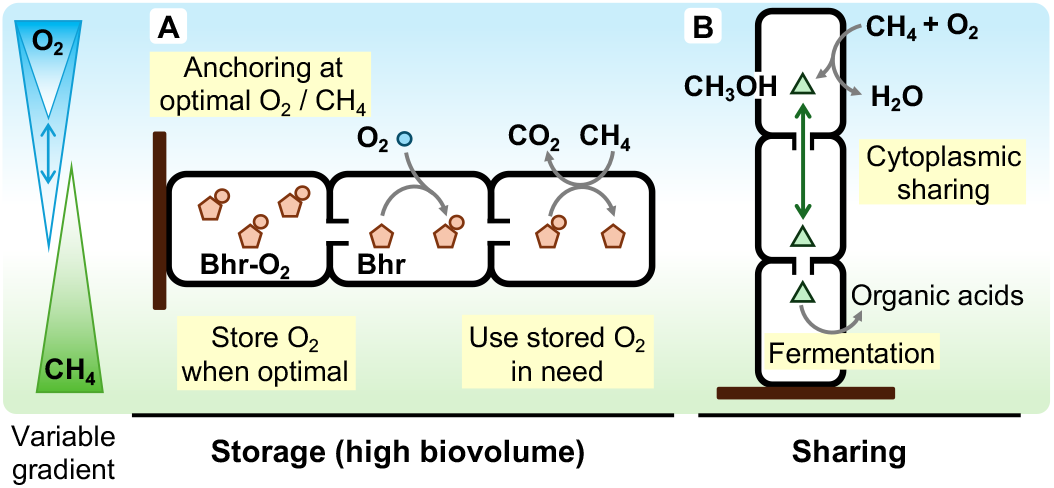
Proposed strategies of filamentous “*Ca*. Allocrenothrix” members to address redox stress. The diagram (not to scale) includes an example O_2_:methane counter-gradient on the left with variability in the steepness of the redox gradient (e.g., O_2_ availability). (A) Cells in filaments may rely on their high biovolumes for storage of metabolic intermediates to withstand transient redox change. (B) In addition, connected cells may share metabolites to remain active across micro-scale redox gradients.

## Conclusion and outlook

Since their description by Cohn, microbiological approaches have gradually illuminated the functional and ecological roles of Crenothrix bacteria in the global environment. Our work represents a critical next step to understanding the basis of the methanotrophic metabolisms of these bacteria, the nature of their cellular interactions, and the adaptations that govern their ecological preferences. Our stable, axenic cultures of “*Ca*. Allocrenothrix methanica” open vast new opportunities for study of the microbial methane cycle, with potential to reveal fundamental ecophysiological drivers of methanotrophy compared to current research that focuses on unicellular and free-living strains. Probing the lifestyle of “*Ca*. Allocrenothrix methanica” in particular could yield important insights into how methanotrophs adapt to seemingly anoxic ecosystems where they are commonly detected [9]. In presenting the isolates in this work, we bring full circle over 150 years of scientific pursuit to discover and describe the microorganisms that drive and impact natural and engineered ecosystems. The pure cultures we present are readily available for study and will allow for the next generation of transformative research in microbial methane metabolism.

### Species description

*Candidatus* Allocrenothrix methanica

#### Etymology

Al.lo.cre.no.thrix. Gr. masc. pron. *allos*, other, another, different; Gr. fem. n. *krênê*, a fountain, spring; Gr. fem. n. *thrix*, hair; N.L. fem. n. *Allocrenothrix*, the other Crenothrix. Me.tha.ni.ca. N.L. neut. n. *methanum*, methane; L. fem. adj. suff. -*ica*, pertaining to; N.L. fem. adj. *methanica*, pertaining to methane.

#### Locality

Isolated from iron-rich wetland sediments collected from Oze National Park, Japan.

#### Diagnosis

Aerobic methane oxidizing-bacterium of the “lacustrine Crenothrix” clade. Grows in linear, rarely branched filaments of cells around 1.0-1.5 µm wide and 1.9-5.0 µm long with no production of gonidia. Cells are connected by conjugation-like structures. There are two strains AF98 and MI19235 for which closed genomes are available (GenBank accession numbers AP041007-AP041015 and AP041016-AP04102, respectively), with AF98 being the proposed type strain.

## Acknowledgements

Gas flux measurements for this project were conducted at Oze National Park as part of the Monitoring Sites 1000 Project of the Ministry of the Environment (Japan). We thank Bernhard Fuchs, Cecilia Wigand, Jörg Wulf, Andreas Ellrott, and Kyoko Kubo for technical assistance with CARD-FISH and LSM image analysis. JMT acknowledges the Young Research Fellow program at the Japan Agency for Marine-Earth Science and Technology.

This research was supported by a Young Scientist Grant from the Institute for Fermentation, Osaka (Y-2024-2-022), a KAKENHI Grant-in-Aid for Scientific Research for Early-Career Scientists from the Japan Society for the Promotion of Science (24K20919), and Joint Research Grants from the Institute of Low Temperature Science at Hokkaido University, Japan (22G006, 23K003, and 24G014). RA was funded by the Max Planck Society.

## Author contributions

K.U., J.M.T., Y.T., and M.F. designed research; K.U., J.M.T., S.N. and M.F. performed field work; K.U. and M.F. performed laboratory experiments; R.A. and M.F. provided key laboratory resources; K.U. and J.M.T. performed data analysis; K.U. and J.M.T. wrote the paper with comments from all authors.

## Data availability

The complete genomes of strains AF98 and MI19235 are available at the DNA Data Bank of Japan (DDBJ) at nucleotide accessions AP041007-AP041015 and AP041016-AP041021, respectively. Raw short (DNBSEQ) and long (Nanopore) DNA sequence reads are available from DDBJ accessions [will be added before publication]. In addition, raw 16S rRNA gene amplicon data (Illumina) for pure culture samples of strains AF98 and MI19235 are available at DDBJ accessions [will be added before publication], and raw 16S rRNA gene amplicon data (Illumina) for wetland samples are available at DDBJ accessions [will be added before publication].

## Supplementary file descriptions

**Supplementary Information** This PDF file includes Figs. S1-8, Tables S1-2, Note S1, Supplementary Methods, and Supplementary References.

**Supplementary Video 1** Time-lapse video of growth of filamentous methanotrophs from 13 h to 14 h incubation time. Strain AF98 is shown under phase-contrast microscopy at 450x speed. The scale bar represents 100 µm.

**Supplementary Video 2** Time-lapse video of growth of filamentous methanotrophs from 35 h to 36 h incubation time. Strain AF98 is shown under phase-contrast microscopy at 450x speed. The scale bar represents 100 µm.

**Supplementary Video 3** Time-lapse video of growth of filamentous methanotrophs from 47 h to 48 h incubation time. Strain AF98 is shown under phase-contrast microscopy at 450x speed. The scale bar represents 100 µm. Interacting filament tips are visible on the bottom left of the video.

**Supplementary Data 1** Information on genomes and gene loci for comparative genomics analyses. The Excel file summarizes assembly statistics for genomes analyzed in this study, locus information for genes and pathways displayed in Fig. 3A, and locus information for key genes identified in the pan-core analyses shown in Fig. S4.

**Supplementary Data 2** Environmental amplicon and physicochemical data. The Excel file contains an amplicon sequence variant (ASV) table summarizing sequence count data for the sampled wetland sites, information about the phylogeny-based classification of *Methylococcales*-associated ASVs, metadata including physicochemical measurements for each site, and raw gas flux measurement values for each site.

**Supplementary Data 3** Environment attribution for global survey results. The Excel file summarizes how metagenome “taxon” names were grouped into broader environment classifications for metagenome hits obtained from the Sandpiper database.

