## Supplementary Information for "Isolation of Crenothrix bacteria reveals the distinct ecophysiologies of filamentous methanotrophs and adaptations to redox stress"

**This PDF file includes:**

- Figs. S1-S8
- Tables S1-S2
- Note S1
- Supplementary Methods
- Supplementary References

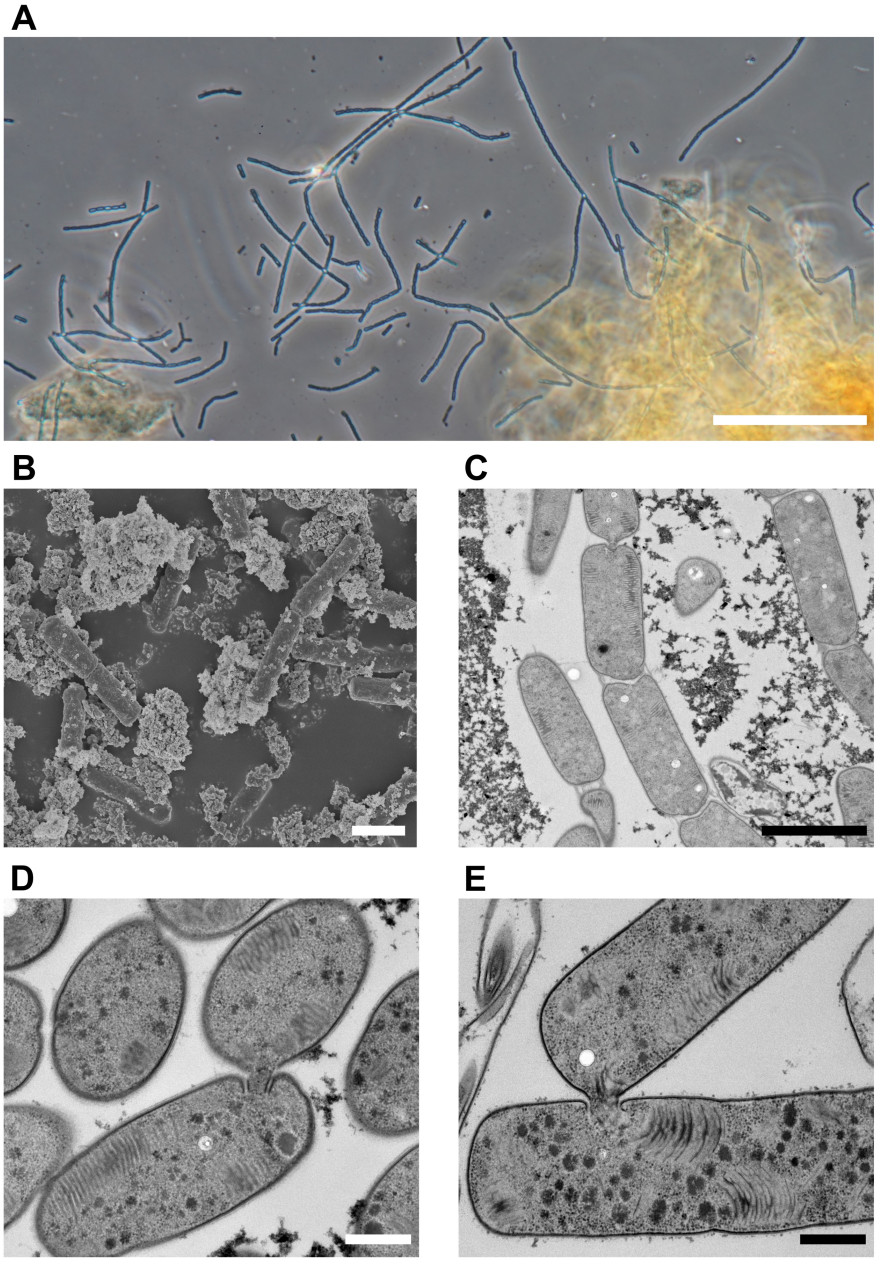


**Fig. S1** Additional microscopy images of filamentous methanotrophs. For all images, biomass was grown under micro-oxic conditions with methane and ferric iron particles. (A) Phase contrast microscopy image of strain MI19235. (B) Scanning electron microscopy image of filaments of strain MI19235. (C) Transmission electron microscopy (TEM) image of strain MI19235 showing internal cellular structures and conjugations between cells. (D,E) TEM images of side-to-side cellular conjugations for strain AF98 (D) and MI19235 (E). Scale bars represent 50 µm (A), 2 µm (B-C) and 0.5 µm (D-E).

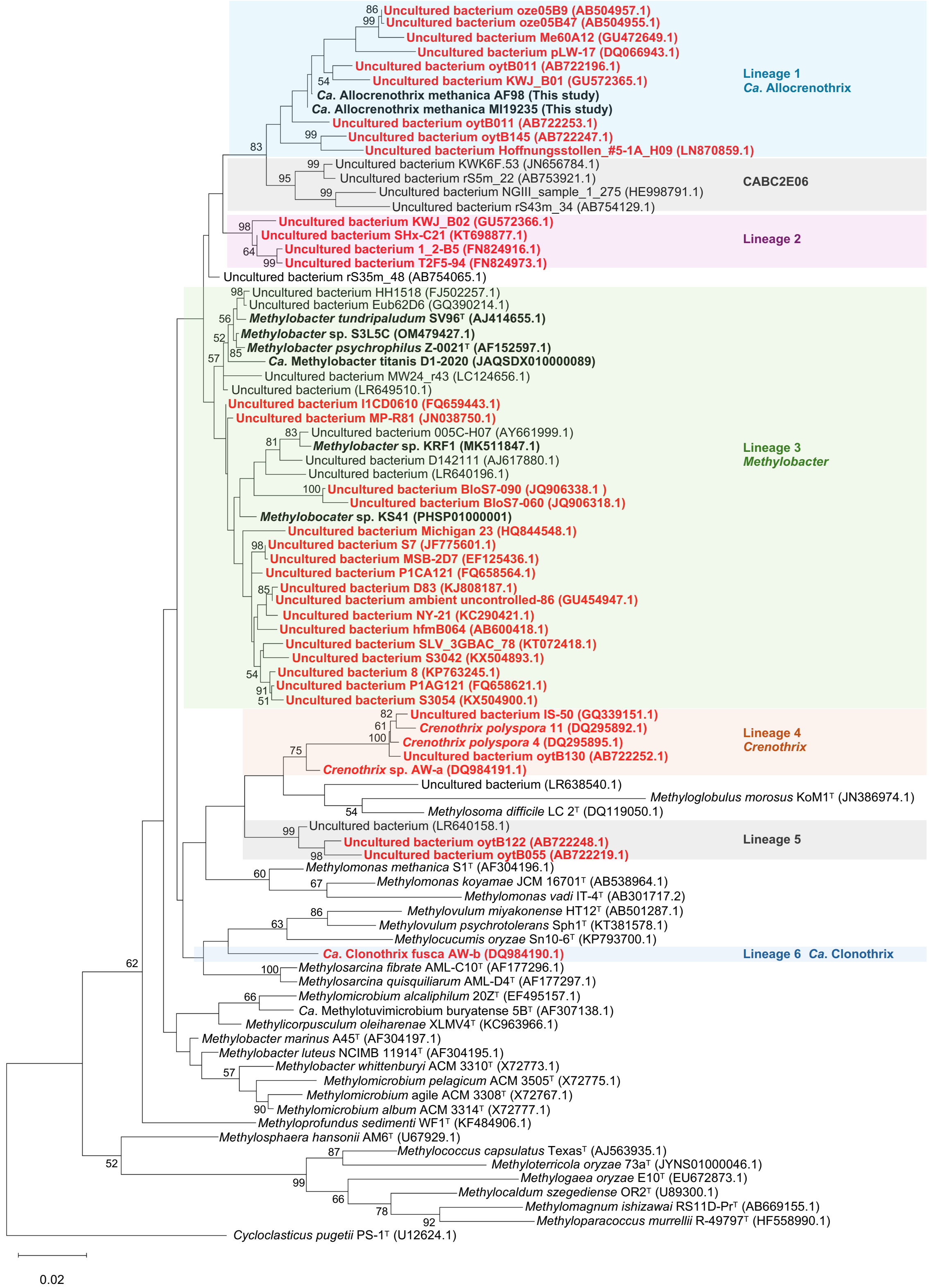


**Fig. S2** Phylogenetic placement of environmental *Methylococcaceae* sequences. The 16S rRNA gene phylogeny includes sequences classified to the *Crenothrix* genus in the Silva database (release 138.2), shown in red, alongside pure culture reference sequences and sequences from this study. Monophyletic sub-lineages are highlighted. Bootstrap values of 50/100 or higher, based on 1000 bootstrap iterations, are shown.

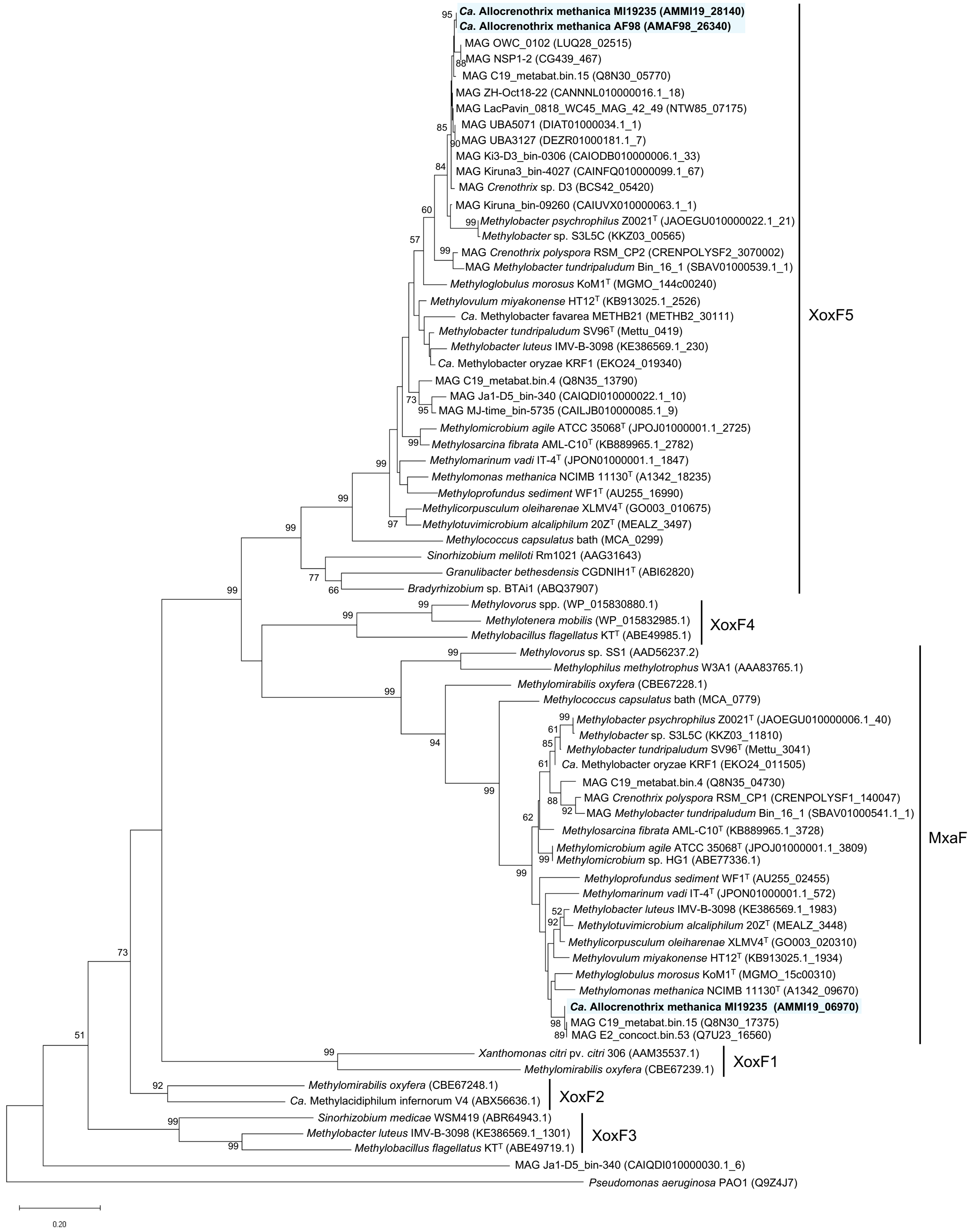


**Fig. S3** Phylogeny of methanol dehydrogenases. The maximum likelihood amino acid phylogeny shows lanthanide-dependent (XoxF) and calcium-dependent (MxaF) dehydrogenases including those detected in strains AF98 and MI19235 (highlighted in blue). Bootstrap values of 50/100 or higher, based on 100 bootstrap iterations, are shown.

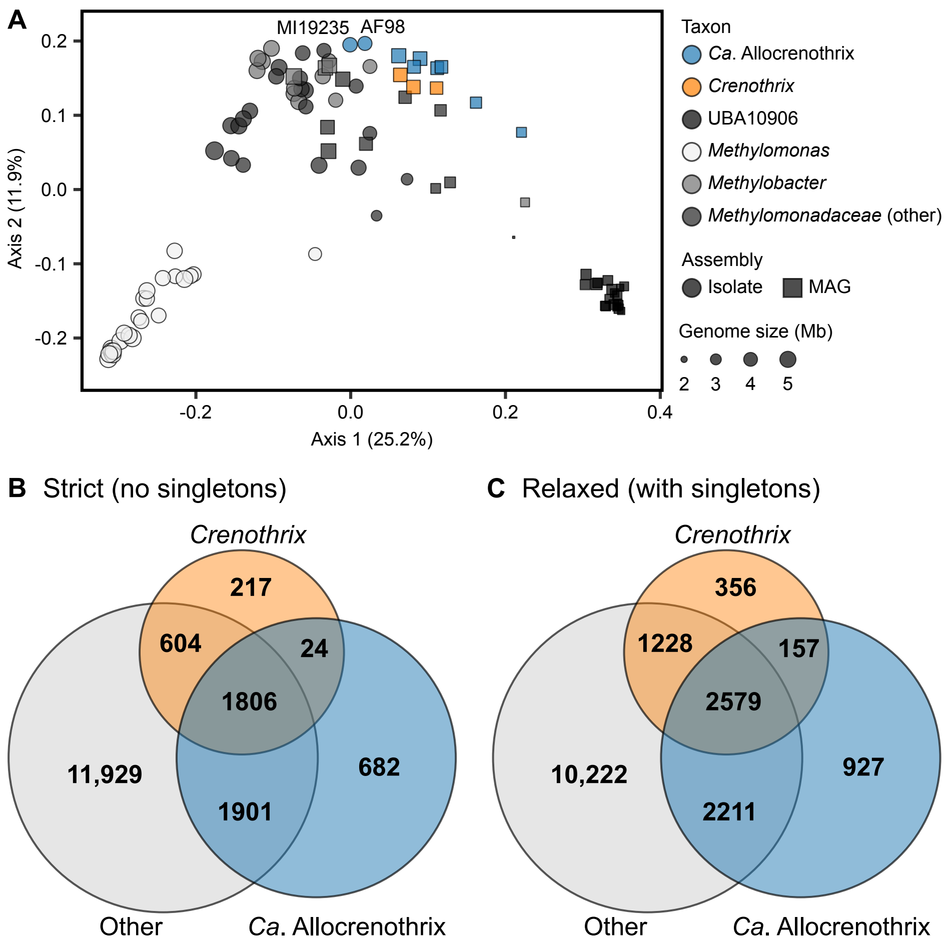


**Fig. S4** Pan-genome analyses of *Methylococcaceae* members. (A) Ordination comparing the genome contents of an expanded set (compared to Fig. 3B) of 99 *Methylococcaceae* members based on counts of orthologous gene groups in genomes. Members of the *Methylomonadaceae* family of the GTDB, which corresponds to the *Methylococcaceae* lineage containing Crenothrix bacteria in NCBI-based taxonomy, were used for analysis (see Supplementary Methods). (B,C) Venn diagrams summarizing pan-core analyses of "*Ca.* Allocrenothrix" members, classical *Crenothrix* members, and other *Methylococcaceae* members. Results are derived from the genome set used in **a** except MAGs were omitted if not classified to "*Ca.* Allocrenothrix" or *Crenothrix*. Both strict (B) and relaxed (C) estimates of core and unique gene counts are shown (see Supplementary Methods for gene counting rules).

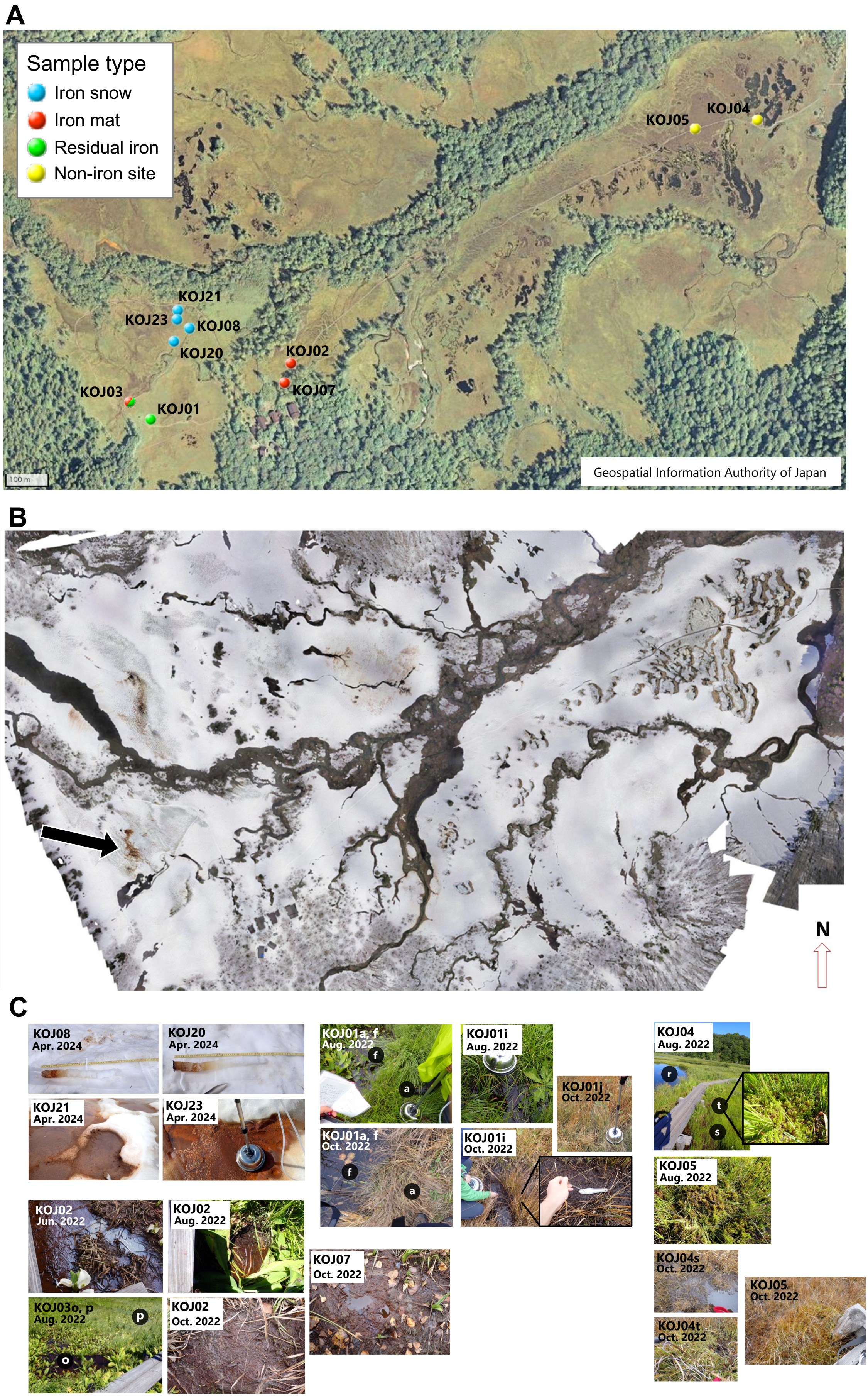


60 **Fig. S5** See next page for caption.

**Fig. S5** Sampling sites for cultivation and gas flux measures. **(A)** Aerial image of the wetland site. Sampling sites are highlighted based on the sample types obtained from each site. (B) Drone image of the same area shown in **a** in winter. An example region with surficial iron snow/mat material is indicated with an arrow. (C) Representative images of selected sites that were sampled for cultivation, microbial community analysis, and gas flux measurements. Sites are organized by their classification as iron snow (top left, four images), iron mats (bottom left, five images), iron residue zones (top center, six images including subset), and non-iron sites (right, six images including subset). Within iron snow sites, the top two images (for KOJ08 and KOJ20) show snow cores, whereas the bottom two images (for KOJ21 and KOJ23) show surficial iron snow. Aerial image (A) is sourced from the Geospatial Information Authority of Japan (https://maps.gsi.go.jp/). Drone image (B) was captured by the authors.

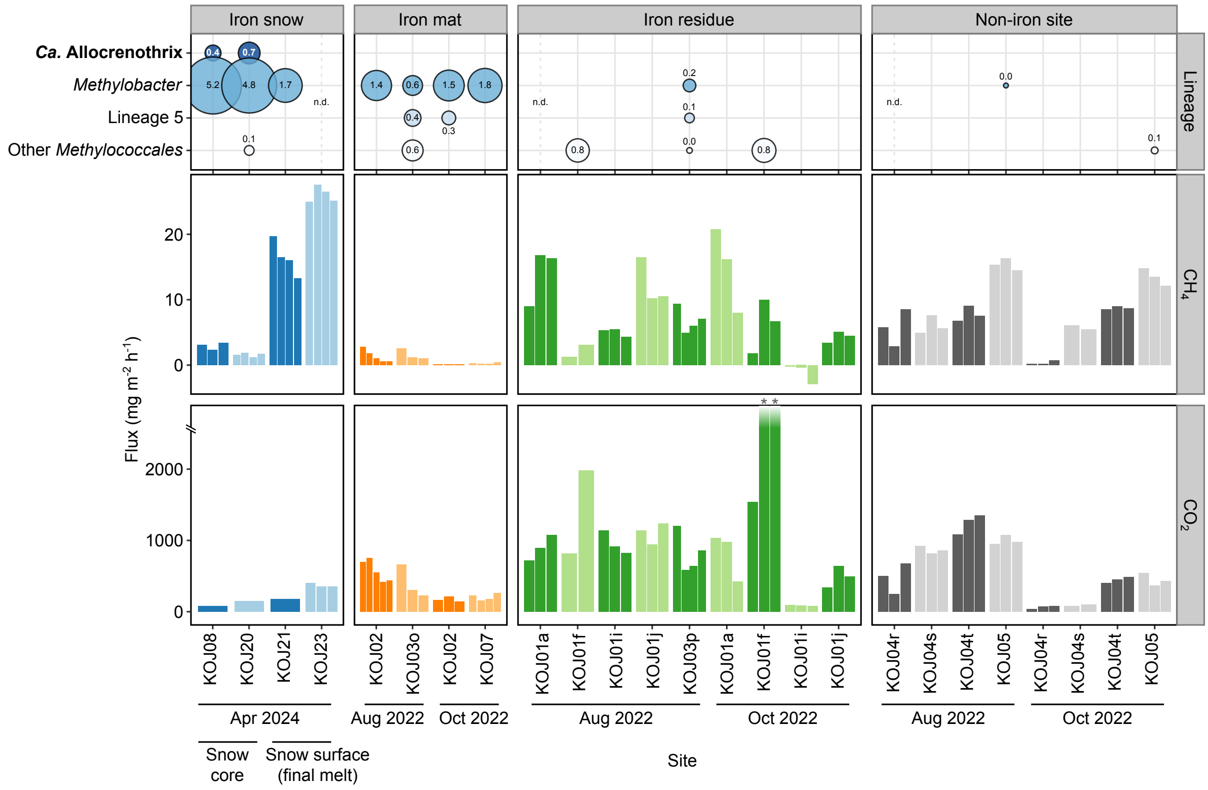


**Fig. S6** Gas fluxes and potential methanotroph communities from wetland samples. The bubble plot shows the relative abundances of potentially methanotrophic taxa, belonging to the *Methylococcales* order, across the analyzed wetland sites based on 16S rRNA gene amplicon data. Amplicon sequence variants (ASVs) with at least 0.3% relative abundance in any sample are shown and are collapsed by lineage based on manual phylogenetic placement (Supplementary Methods, Fig. S2). For snow core KOJ20, the 0-6 cm slice is shown (see Fig. 4B and Supplementary Data 2 for other depth slices). Bar plots show the measured fluxes of methane (top) and carbon dioxide gas (bottom) from the wetland sites, with technical replicate measures shown as adjacent bars. The two carbon dioxide flux values for site KOJ01f (October 2022) that exceed the scale area, indicated by asterisks, are 8,399 and 5,764 mg $m^{-2}$ $h^{-1}$. Measurement sites are shown in Fig. S5. Abbreviations: n.d., no data.

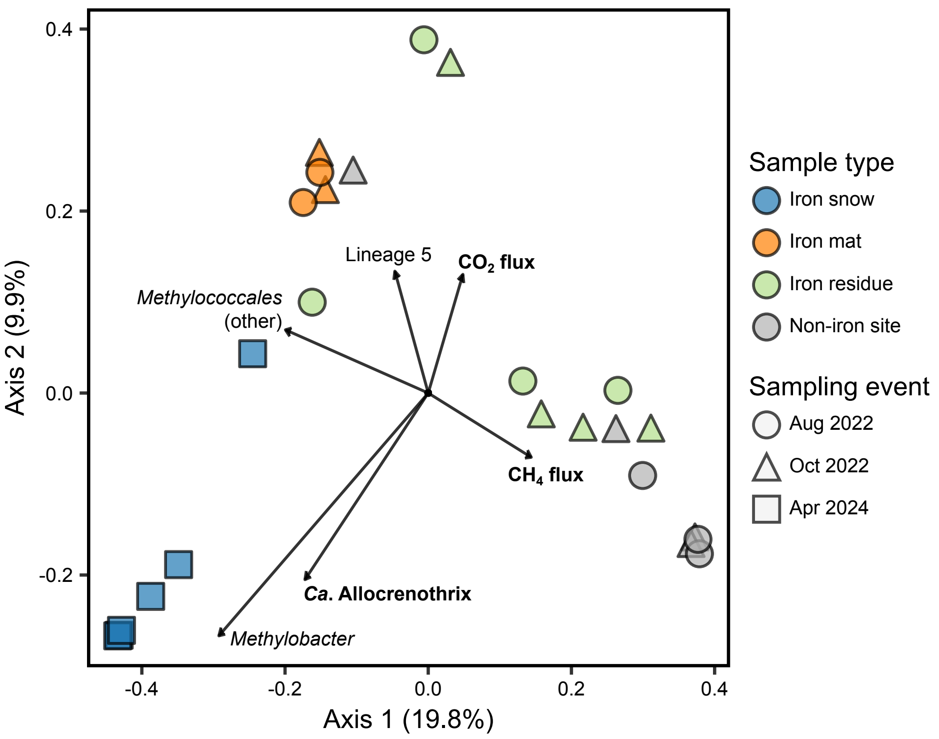


**Fig. S7** Ordination of the microbial community structure of wetland samples. The distance biplot shows the relative magnitude of correlation of each metadata factor (indicated by an arrow) with microbial community samples (indicated as points). Samples are separated based on Bray-Curtis dissimilarity via principal coordinate analysis (PCoA). The same samples are shown as in Fig. S6 except that additional slices of KOJ20 snow cores are included (see Fig. 4A and Supplementary Data 2). The proportion of variance in the microbial community data explained by each axis is indicated beside the axis label. Potential methanotroph taxa included in the biplot are the same as those shown in Fig. S6.

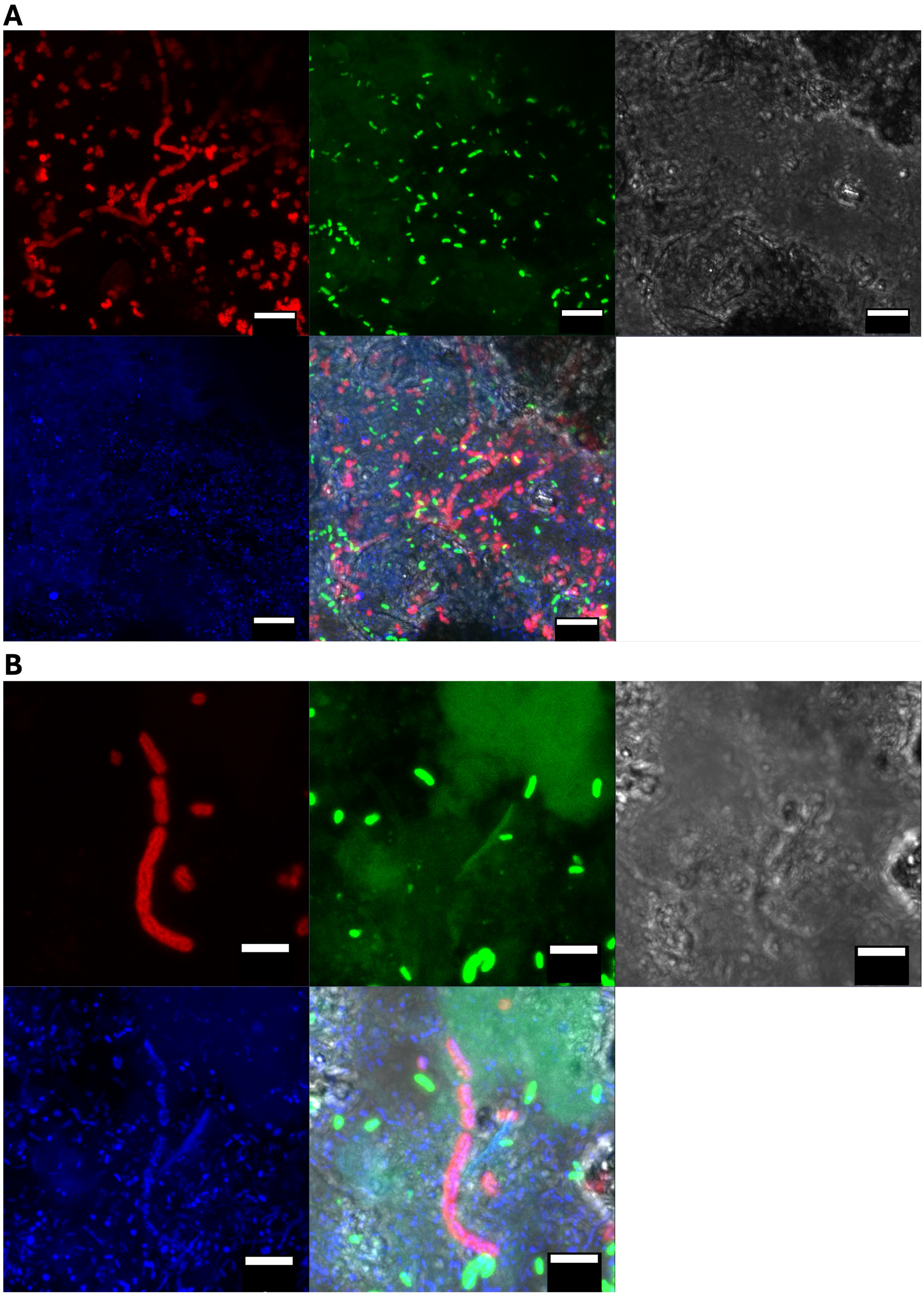


**Fig. S8** Additional CARD-FISH microscopy images from iron-rich snow core samples. (A,B) Images from sites KOJ20 (A; core slice 6-12 cm above surface) and KOJ08 (B; core slice 4.5-11 cm above surface). In both (A) and (B), the order of the image tiles (left to right, top to bottom) is: Mγ669 probe (red), GEO3 A-C probes (green), differential interference contrast microscopy (DIC; grey), DAPI staining (blue), overlay. (The bottom right tile is intentionally left empty.) Scale bars represent 10 μm (A) and 5 μm (B).

**Table S1** Composition of media used for cultivation of filamentous methanotrophs.

| Constituent | Oz-dNMS | Sv-dNMS | PSVn | PSVal |
|---|---|---|---|---|
| Basal constituents | | | | |
| $KNO_3$ | 0.2 g | 0.2 g | 0.2 g | - |
| $NH_4Cl$ | - | - | - | 0.1 g |
| $MgSO_4 \cdot 7H_2O$ | 0.2 g | 0.2 g | 0.2 g | 0.2 g |
| $CaCl_2 \cdot 2H_2O$ | 0.2 g | 0.2 g | 0.2 g | 0.2 g |
| $KH_2PO_4$ | - | 0.02 g | 0.02 g | 0.02 g |
| MES/NaOH | 4 mM | 4 mM | - | - |
| PIPES/NaOH | - | - | 4 mM | 4 mM |
| Trace element solution† | 1 mL | 1 mL | 1 mL | 1 mL |
| Selenite-tungstate solution† | 1 mL | 1 mL | 1 mL | 1 mL |
| 100 µM $La(NO_3)_3$ solution | - | - | (1 mL)* | 1 mL |
| Vitamin mixture solution† | 1 mL | 1 mL | 1 mL | 1 mL |
| 35 mM phosphate buffer (pH 5.6) | 2 mL | - | - | - |
| Headspace‡ | A | A/B | A/C | D |

* Optional medium component.

† Composition is described in the Supplementary Methods.

‡ Gas composition: A: 47.5:2.5:25:25 $N_2$:$CO_2$:$CH_4$:air (1 atm); B: 71.3:3.8:25 $N_2$:$CO_2$:$CH_4$ (1 atm); C: 63.3:3.3:16.7:16.7 $N_2$:$CO_2$:$CH_4$:air (1.5 atm); D: 76:4:20 $N_2$:$CO_2$:$CH_4$ (1.25 atm).

The composition per litre is shown for the four medium variants developed in this work. The Oz-dNMS and Sv-dNMS variants were used for early cultivation work, whereas the PSVn and PSVal variants represent more highly optimized medium variants. See the Supplementary Methods for additional details.

**Table S2** Summary statistics for (bacterio)hemerythrin and partial denitrification genes.

| | (Bacterio)hemerythrin (median count per genome) | | | Partial denitrification (% occurrence in genomes) | | |
|---|---|---|---|---|---|---|
| **Genome group** | **1*-Bhr[†]** | **2-Hr[‡]** | **3-Hr** | **1** | **2** | **3** |
| "*Ca.* Allocrenothrix" | 5 | 8 | 8 | 81.3 | 66.7 | 66.7 |
| *Crenothrix* | 3 | 2 | 2 | 0.0 | 0.0 | 0.0 |
| UBA10906 | 3 | 3 | 3 | 50.0 | 31.2 | 31.6 |
| *Methylococcaceae* (other)[§] | 3 | 5 | 3 | 62.5 | 58.8 | 35.0 |
| *Methylococcaceae* (GTDB)[§] | - | - | 4 | - | - | 22.7 |
| *Methylococcales* (other) | - | - | 1 | - | - | 9.8 |

*Genome groups used for statistics: **1**: *Methylococcaceae* genome set (37 genomes) in Fig. 3a;
**2**: *Methylococcaceae* genome set (99 genomes) in Supplementary Fig. 4a;
**3**: Full set of 313 *Methylococcales* genomes (see Supplementary Methods).

[†] Bhr (bacteriohemerythrin) members are those with hemerythrin domains that passed manual screening as potential bacteriohemerythrin protein candidates based on CD-search results (see Methods). CD-search based screening was only performed for genome subset 1.

[‡] Hr (hemerythrin domain-containing) members are those with hemerythrin domains identified by eggnog-mapper (see Supplementary Methods).

[§] The GTDB divides classical *Methylococcaceae* members into multiple taxonomic groups. Here, *Methylococcaceae* (other) refers to members of the *Methylomonadaceae* family in the GTDB outside of "*Ca.* Allocrenothrix", *Crenothrix*, and UBA10906 members. *Methylococcaceae* (GTDB) refers to a separate family called *Methylococcaceae* in the GTDB that does not include Crenothrix bacteria. Members of both of these families in the GTDB are generally classified together as members of *Methylococcaceae* based on NCBI taxonomy.

The table summarizes the occurrence of bacteriohemerythrin gene candidates and genes predicted to encode hemerythrin domains within the three analyzed genome sets. Occurrence of genes encoding the partial denitrification pathway from nitrite to nitrous oxide are also shown. Gene targets are described in detail in the Methods and Supplementary Methods.

### Supplementary Notes

*Note S1*

**Additional comparative genomic analyses of Methylococcaceae members.** We further analyzed the genomes of strains AF98 and MI19235 and compared them to the set of 37 genomes and MAGs classified to *Methylococcaceae* family (Fig. 3A). Although we found that strains AF98 and MI19235 lacked genes for soluble methane monooxygenases (*mmo* genes), some MAGs of uncultured "*Ca.* Allocrenothrix" members possessed *mmoXYBZDC* and lacked *pmoCAB*. In general, "*Ca.* Allocrenothrix" members encoded methanol dehydrogenase genes and genes for subsequent metabolism of methanol into carbon dioxide or biomass, including genes for the $H_4$MPT and RuMP pathways [50, 74], supporting that "*Ca.* Allocrenothrix" members utilize the classical pathway of aerobic methane oxidation (Fig. 3A). Lanthanide-dependent methanol dehydrogenase gene were commonly encoded by "*Ca.* Allocrenothrix" genomes, and this leads us to speculate lanthanide dependence could be one reason for previous failures in cultivating "*Ca.* Allocrenothrix" members given that most methanotroph medium has historically lacked such trace metals [55]. We noted that none of the "*Ca.* Allocrenothrix" genomes or MAGs encoded a *rbcL* form I RuBisCO large subunit gene associated with carbon fixation, although this gene is known to be present in the genome of the model methanotroph *Methylococcus capsulatus* Bath.

In the genomes of strains AF98 and MI19235, we detected the complete gene pathway for fermentative production of acetate from pyruvate, as was described for *Methylobacter* sp. S3L5C (Fig. 3C) [50]. In addition, all genes were detected for production of propionate from pyruvate except the propionyl-CoA:succinyl-CoA transferase *scpC*, and we also detected *hox* genes for [NiFe]-hydrogenases that could potentially be linked to fermentative hydrogen gas production [50]. MAGs of other "*Ca.* Allocrenothrix" members similarly included genes for fermentation to propionate from pyruvate but lacked *scpC*. Beyond this, we searched for genes implicated in the fermentative pathways of *Methylotuvimicrobium alcaliphilum* 20Z in "*Ca.* Allocrenothrix" MAGs and the AF98/MI19235 genomes, but many key genes were lacking, including *ldh*, *sfcA, fumC* and *pta* genes encoding lactate/malate dehydrogenase, malate dehydrogenase, fumarate hydratase and phosphate acetyltransferase, respectively [11]. Within all analyzed *Methylococcaceae* genomes, only the *Methylobacter psychrophilus* Z0021$^{T}$ and *Methyloprofundus sediment* WF1$^{T}$ genomes possess the complete gene sets for fermentation described for *Methylobacter* sp. S3L5C and *Methylotuvimicrobium alcaliphilum* 20Z, respectively, implying these exact fermentation pathways may not be broadly shared among *Methylococcaceae* members.

We also detected homologs of *cyc2* and *cyc1*, which have been implicated in iron metabolism and extracellular electron transfer [75], in the genomes of strains AF98 and MI19235. The detected homologs of *cyc2* placed into "cluster 2", a clade of *cyc2* linked to known iron-oxidizing bacteria such as *Acidithiobacillus ferrooxidans* [76], based on analysis by FeGenie [29]. Similar *cyc2* gene homologs were not detected in "*Ca.* Allocrenothrix" MAGs but were encoded by several other *Methylococcaceae* genomes (Fig. 3A). The role of *cyc2* among methanotrophs remains unclear.

Many bacteria rely on siderophores to obtain insoluble ferric iron. Given the potential use of iron in bacteriohemerythrins by strains AF98 and MI19235, we checked the genomes of both strains for genes involved in siderophore biosynthesis. In both genomes, we detected a putative gene cluster involved in siderophore biosynthesis and confirmed the identity of the included genes by manual homology searches via BLASTP [77]. Along with several genes encoding non-ribosomal peptide synthetases and a TonB-dependent receptor protein, these clusters included the *pvdS* (AF98: locus tag AMAF98_30710, MI19235: locus tag AMMI19_12660) and *pvdE* (AF98: locus tag AMAF98_30700, MI19235: locus tag AMMI19_12670) genes involved in biosynthesis of pyoverdine siderophores in *Pseudomonas aeruginosa* PAO1 [78], and the *fpvG* gene (AF98: locus tag AMAF98_30780, MI19235: locus tag AMMI19_12570) involved in reduction of chelated ferric iron for release from siderophores [79]. Detection of these genes implies that both strains may be able to uptake insoluble ferric iron by chelation with siderophores.

For nitrogen metabolism, strain AF98 encodes the nitrite reductase *nirK* [63]. Although lacking *nirK*, strain MI19235 encodes a putative homolog of the alternative nitrite reductase *nirS* [63], which implies that both strains may be capable of nitrite reduction.

On the plasmids of strains AF98 and MI19235, we found a *nifH* gene homolog potentially encoding a nitrogenase (plasmids AP041010 and AP041019 for AF98 and MI19235, respectively). We also identified *vir* genes involved in type IV secretion systems (MI19235: *virB1*-*virB11*; AF98: same but lacking *virB3* and *virB6*) on plasmids AP041009/AP041011 and AP041017 for AF98 and MI19235, respectively. *vir* genes were also detected in the genomes of select other "*Ca*. Allocrenothrix" members and another *Methylococcaceae* strain (Fig. 3A). One bacteriohemerythrin-like gene (locus tag AMMI19_37490) of strain MI19235 was also found on the plasmid AP041018, although the other potential bacteriohemerythrin genes of this strain, along with all potential bacteriohemerythrin genes of strain AF98, were found on chromosomes.

In all comparative genomics analyses, we note that aside from our complete genome sequences, the "*Ca*. Allocrenothrix" and *Crenothrix* genera are represented by uncultured MAGs with moderate to severe fragmentation (Fig. 3A), allowing for potential mis-binning of contigs. Yet our detection of genes related to those encoded by the complete genomes of strains AF98 and MI19235 in other uncultured "*Ca*. Allocrenothrix" members supports that multiple genomic features of strains AF98 and MI19235 may be common throughout the "*Ca*. Allocrenothrix" genus.

### Supplementary Methods

*Cultivation medium*

Medium used during cultivation was based on diluted nitrate or ammonium mineral salts medium and was iteratively refined during our cultivation experiments (Table S1) [80–82]. The variants of the medium we ultimately used for isolation and maintenance of the strains, named PSVn and PSVal, included 0.2 g $L^{-1}$ magnesium sulfate heptahydrate, 0.2 g $L^{-1}$ calcium chloride dihydrate, and 0.02 g $L^{-1}$ potassium phosphate monobasic, along with 0.2 g $L^{-1}$ potassium nitrate (for PSVn) or 0.1 g $L^{-1}$ ammonium chloride (for PSVal). Piperazine-N,N-bis(2-ethanesulfonic acid) (PIPES) was added as an organic pH buffer to a final concentration of 1.2 g $L^{-1}$ (4 mM), and the final pH of the medium was adjusted to 6. In addition, 1 mL $L^{-1}$ each of a previously described non-chelated trace element solution [83] and selenite-tungstate solution [83] were added, along with 1 mL $L^{-1}$ of a vitamin solution, modified from Wolin [84], which was composed of the following (in mg $L^{-1}$): biotin, 2; folic acid, 2; pyridoxine hydrochloride, 10; thiamine HCl, 5; riboflavin, 5; nicotinic acid, 5; calcium pantothenate, 5; vitamin B12, 0.1; p-aminobenzoic acid, 5; DL-alpha-lipoic acid, 5. For PSVal medium, 1 mL $L^{-1}$ of 100 µM lanthanum(III) nitrate solution was always added, whereas this was an optional supplement to PSVn medium (and was not added for the experiments based on PSVn medium described in this work unless otherwise indicated).

Medium was prepared under anoxic conditions with dinitrogen and carbon dioxide gas in the headspace and sterilized as in [83]. Sterile methane and/or air were then added to the headspace by injection. Unless otherwise specified, the final gas composition of the medium was 47.5:2.5:25:25 of dinitrogen:carbon dioxide:methane:air, at 1 atm final pressure, for PSVn, and was 76:4:20 of dinitrogen:carbon dioxide:methane, at 1.25 atm final pressure, for PSVal. In addition, poorly crystalline ferric oxide particles were prepared by neutralizing a ferric chloride solution with NaOH and washing with water to desalt, as described previously [85], for optional addition into the medium.

During our initial cultivation efforts, early variants of these media called Oz-dNMS and Sv-dNMS were used (see Table S1). Compared to PSVn medium, these media relied on 2-(N-morpholino)ethanesulfonic acid (MES) as an organic pH buffer (4 mM concentration), and the lanthanum solution was omitted. In addition, for Oz-dNMS, the potassium phosphate monobasic was substituted for a 35 mM phosphate solution (pH=5.6), which was added at 2 mL $L^{-1}$ to the medium.

*Enrichment and isolation of strains*

In June 2022, Oz-dNMS medium was used with a micro-oxic (approximately 5% $O_2$) headspace containing 25% methane gas, and poorly crystalline ferric oxide particles were added to a final iron concentration of 10 mM. After inoculation, serial dilution in medium was immediately performed before incubation at 15°C in the dark. Inoculation and incubation in August 2022 was done similarly, except Sv-dNMS medium was used with an anoxic headspace, and serial dilution was not performed. Strains MI19235 and AF98 were isolated from June and August cultures, respectively, following the methods below.

After initial growth, the MI19235 enrichment culture was subcultured several times in Oz-dNMS medium with 10 mM poorly crystalline ferric oxide particles, followed by several subcultures in Oz-dNMS or Sv-dNMS medium without ferric oxide particles. To remove small contaminant cells, culture material was passed through a 0.8 µm centrifugal filter unit (Centrex centrifugal microfilter-sterile; GVS, Bologna, Italy) by centrifugation at 1500 x g for 10 min. Part of the filter was then inoculated into SV-dNMS medium, and serial dilution was performed.

To further refine the MI19235 enrichment culture, cells were sorted using glass micropipettes under an inverted microscope. First, the culture was passed through a 0.8 µm filter unit as above. Medium filtered via a 0.22 µm filter, to remove small debris that could obstruct observations, was used to resuspend cells from the 0.8 µm filter, and the suspension was added to a glass dish for processing. Ultra-thin glass micropipettes were prepared by stretching and bending glass pipettes (G-1; Narishige, Tokyo Japan or B100-75-10; Sutter instrument, Novato, CA) under flame, either manually or using a puller (P-97; Sutter instrument) and micro forge (MF-900; Narishige), followed by sterilization at 180-190°C for 1 h. Single filaments or bunches of filaments were then sorted from the prepared culture material via the glass micropipettes using a microinjector (IA-1; S, Tokyo, Japan) operated with a micromanipulator (3-man; S) under phase-contrast microscopy via an inverted microscope (IX71; Evident, Tokyo, Japan). Sorted cells were immediately inoculated into fresh Sv-dNMS medium.

After multiple additional rounds of subcultures in Sv-dNMS or PVSn medium under micro-oxic conditions (approximately 3.5-5% $O_2$) with methane, treatment via 0.8 µm filtration, micropipette-based cell sorting (with the pre-filtration step omitted; including 0.1 mM poorly crystalline iron oxide), and serial dilution, we obtained a pure culture of strain MI19235. A total of 48 subcultures were performed from initial enrichment to isolation.

The initial subculture of the strain AF98 enrichment (derived from an August 2022 sample) was performed in anoxic PSVal medium containing 0.1 µM lanthanum(III) nitrate and 10 mM poorly crystalline ferric oxide with a headspace of 76:4:20 dinitrogen:carbon dioxide: methane (1.25 atm pressure). Subsequently, a subculture was performed in PSVn medium without lanthanum or poorly crystalline ferric oxides under micro-oxic conditions with methane. Cell sorting via micropipette was then performed as described above, and sorted cells were inoculated into PSVn medium using the same conditions as the previous subculture. In the subsequent subculture, serial dilution was performed in PSVn medium with the same conditions as above. Strain AF98 was successfully isolated after this subculture, which was the fifth total subculture after its initial enrichment.

Following isolation, both strains were generally maintained in PSVn medium under micro-oxic conditions with methane. To cultures of strains AF98, 1-10 mM of poorly crystalline ferric oxide and 0.1 µM lanthanum(III) nitrate were typically added to the medium.

*Additional details for amplicon and genome analyses of cultures*

Amplicon sequencing to confirm the purity of cultures was performed by the Bioengineering Lab Co. (Sagamihara, Japan). The V4 region of the 16S rRNA gene was amplified via polymerase chain reaction (PCR) using LongAmp Hot Start Taq 2x master mix (New England Biolabs, Ipswich, MA) with the PCR primers 515f (5'-GTG CCA GCM GCC GCG GTA A-3') and 806r (5'-GGA CTA CHV GGG TWT CTA AT-3') attached to linking adaptors[86]. Sequence libraries were prepared by attaching amplicons, purified using VAHTS DNA Clean Beads (Vazyme; Nanjing, China), to sequencing adaptors with index sequences via a second round of PCR using KOD FX Neo polymerase (TOYOBO, Osaka, Japan). Libraries were sequenced using the MiSeq platform (2x300 bp) with Miseq Regent Kit v3 (Illumina, San Diego, CA).

Primers of amplicon reads were trimmed by FASTX-Toolkit version 0.0.14 (https://github.com/agordon/fastx_toolkit), and reads were quality controlled by sickle version 1.33 (https://github.com/najoshi/sickle). Denoising, clustering and removing chimeric sequences were performed by DADA2 via QIIME2 version 2024.2 [24, 25]. The detection limit of sequencing was calculated by dividing the minimum allowable sequence count (two reads) by the total post-denoising read counts of a sample, and low-abundance sequence variants with one mismatch to the dominant variant were considered part of the same sequence cluster.

Long-read genome sequencing is described in the Methods. For short-read genome sequencing by the Bioengineering Lab Co., libraries were prepared using the MGIEasy FS DNA Library Prep Set (MGI Tech, Shenzen, China) and circulated using the MGIEasy Circularization Kit (MGI Tech). DNA nanoballs were produced using DNB Rapid Make Reagent Kit (MGI Tech) and sequenced with the DNBSEQ-T7 platform (2x150 bp reads).

Initial quality control was performed for long-read genome data to remove control strands via CLEAN version 1.0.2 [87]. Porechop version 0.2.4 (https://github.com/rrwick/Porechop) was also used to remove chimeric sequences of long reads and trim reads adapters. Adaptors were trimmed from short-read genome data using cutadapt version 4.9 [88]. The long Nanopore reads and the short DNBSEQ reads were then used for hybrid genome assembly via the Hybracter pipeline [26], version 0.7.4. Steps in the pipeline included quality control using filtlong version 0.2.1 (https://github.com/rrwick/Filtlong), porechop version 0.5.0, seqkit version 2.5.0 [89], and fastp version 0.23.4 [90]; long-read assembly using Flye version 2.9.4-b1799 [91], with varied subsampling depth between 100-200x, and plasmid assembly using Plassembler version 1.6.2 [92]; sequence polishing using Medaka version 1.8.0 (Oxford Nanopore Technologies; https://github.com/nanoporetech/medaka), Polypolish version 0.6.0 [93], and Pypolca version 0.3.1 [94, 95]; and chromosome reorientation using Dnaapler version 0.8.0 [96].

*Sample preparation for electron microscopy*

For transmission electron microscopy, harvested cells were held between copper plates and rapidly frozen in liquid propane (-175°C). Cells were then fixed by freeze substitution for

48 h at -80°C using a solution of 2% glutaraldehyde / 1% tannic acid in ethanol / 2% distilled water, and samples were then incubated at -20°C for 3 h and 4°C for 3 h. After multiple dehydration treatments in 100% ethanol, samples were incubated in propylene oxide for 30 min twice and immersed in equal volumes of propylene oxide and epoxy resin (Quetol 651; Nisshin EM, Tokyo, Japan) for 3 h. After propylene oxide was volatilized, samples were embedded in epoxy resin (Quetol 651; Nisshin EM) for 48 h at 60°C. Ultrathin sections (70 nm) were then created by slicing samples using an ultramicrotome (Ultracut UCT; Leica Vienna, Austria). Following this, samples were stained with 2% uranyl acetate and lead stain solution (Merk; Darmstadt, Germany) for 15 min and 3 min, respectively, before being used for observation.

For scanning electron microscopy, harvested cells were suspended in a solution of 4% paraformaldehyde, 4% glutaraldehyde, and 0.1 M cacodylate buffer (pH=7.4) and cooled to 4°C, and then cells were fixed in 2% glutaraldehyde in 0.1 M cacodylate buffer (pH=7.4) overnight at 4°C. After washing cells with 0.1 M cacodylate buffer three times, cells were fixed with 2% osmium tetroxide in 0.1 M cacodylate buffer for 90 min at 4°C. Fixed samples were subjected to an ethanol dehydration series (50%, 70%, 90% and 100%), incubated in mixture of ethanol and t-butyl alcohol (1:1) followed by three 30 min incubations in t-butyl alcohol, and then freeze dried. Freeze dried samples were coated with a layer of 30 nm osmium and then used for observation.

*Additional details for comparative genomics analyses*

Construction of the concatenated marker protein tree via GToTree version 1.8.3 [32] for the set of 37 *Methylococcaceae* genomes/MAGs (plus outgroup) was performed as follows. Single copy genes for *Gammaproteobacteria* (172 target genes) were detected using HMMER3 version 3.3.2 [42] among open reading frames predicted by Prodigal version 2.6.3 in the genomes [97]. Of the 172 target genes, 170 genes were detected, and these were aligned and trimmed by Muscle version 5.1 [98] and TrimAl version 1.4.1 [99]. The maximum likelihood tree was then constructed via IQ-TREE version 2.2.5 [100] using the LG+F+I+R5 evolutionary model, as selected by ModelFinder [101], with 1000 ultrafast bootstraps [102].

To complement analyses performed on the set of 37 *Methylococcaceae* genomes, genomes of additional *Methylococcales* members were downloaded and analyzed. All GTDB species representatives (release 226) belonging to the *Methylococcales* order with >80% completeness and <5% contamination based on CheckM2 [103] were initially targeted for analysis. Genomes/MAGs were downloaded using bit version 1.13.2 (https://github.com/AstrobioMike/bit), and genes were predicted using DFAST version 1.3.7[27] with the '--metagenome' flag. Combined with the complete genomes of strains AF98 and MI19235, this resulted in a collection of 313 genomes. OrthoFinder version 3.1.0 [38] was used to identify orthologous gene groups in the genome collection. Singleton orthologous groups (i.e., having only one gene member across the entire set) were excluded from subsequent analyses.

An approximate functional annotation of orthologous gene groups was performed by selecting one gene member per group (from AF98 or MI19235, if possible) and annotating using eggnog-mapper version 2.1.3 [104, 105]. Orthologous groups of hemerythrin domain-containing proteins were identified if the "PFAMs" annotation results from eggnog-mapper included a hit to the "Hemerythrin" family. Median counts of hemerythrin domain-containing genes per genome were then summarized per methanotroph lineage. For denitrification genes, reference sequences for NirK, NirS, and NorB from strain AF98 were queried against the sequence set used for eggnog-mapper annotations using BLASTP version 2.16.0 [77] with an e-value threshold of $10^{-10}$. Orthologous gene group hits were then screened by web BLASTP search [106] against the ClusteredNR database (September 2025) and by manual evaluation of closest matches. A genome was considered to have potential for partial denitrification from nitrite to nitrous oxide if the genome included at least one count of a *nirK* or *nirS* orthologous group as well as at least one count of a *norB* orthologous group. Relative occurrence of the partial denitrification pathway in genomes was then summarized per methanotroph lineage.

To avoid potentially spurious results due to quality of MAGs, the expanded genome set was then subset to all cultured isolates belonging to the *Methylomonadaceae* family of the GTDB (which corresponds to the *Methylococcaceae* lineage containing Crenothrix bacteria in NCBI taxonomy), along with MAGs belonging to the *Crenothrix*, "*Ca.* Allocrenothrix", or UBA10906 genera, the genomes of strains AF98 and MI19235, and highly contiguous MAGs belonging to the *Methylomonadaceae* family with a scaffold L50 score of 10 or less. A total of 99 genomes were obtained in the subset. The above tests for hemerythrin domain-containing proteins and partial denitrification were repeated for this genome subset. In addition, a principal coordinate analysis (PCoA)-based ordination was constructed based on orthologous gene group counts as in the Methods. Lastly, a pan-core analysis was performed. Non-Crenothrix bacterial MAGs were omitted from the analysis to avoid errors due to mis-binning, leaving 67 "*Ca.* Allocrenothrix", *Crenothrix*, and other *Methylococcaceae* genomes. The pan-core analysis was performed using both strict and relaxed gene counting rules. In the strict analysis, orthologous gene groups had to be present in at least two genomes in a lineage to be considered present in that lineage. By comparison, in the relaxed analysis, an orthologous gene group was considered present in a lineage if at least one genome included the orthologous gene group, although the membership threshold for the "other *Methylococcaceae*" lineage was set to be at least two genomes to avoid bias towards this larger genome set. Unique and shared gene counts between the three lineages were then summarized.

*Environmental sample collection details*

Samples for DNA extraction were collected from mat or soil sites by mixing soil/sediment material 1:1 with 2x DNA/RNA Shield (Zymo Research; California, U.S.A.). Select samples (i.e., sub-sites KOJ1f and KOJ4r) were taken from ponds in the wetland, and for these, surface water or surficial sediment mixed with overlying pond water was preserved 1:1 in 2x DNA/RNA Shield similarly. For one water sample, water was passed through a Sterivex filter (Merck; Darmstadt, Germany), and the filter was fixed with 1.8 mL of 1x DNA/RNA Shield. In the spring melt season (for sites KOJ08 and KOJ20), snow cores were taken using an acrylic tube-based sampling device and were then separated by depth. Material from snow cores was collected for DNA extraction by allowing the snow to melt over two days while cool. Melted material was then passed through Sterivex filters (Merck) and preserved with 1x DNA/RNA Shield. For snow surface samples (at sites KOJ21 and OJ23), surficial iron precipitate that had accumulated on the top of the snow was directly mixed 1:1 in 2x DNA/RNA Shield. In addition, for CARD-FISH analyses, core slices from 6-12 cm and 4.5-11 cm above surface was used for fixation for sites KOJ20 and KOJ08, respectively, as described in the Methods.

Air temperatures at sampling sites were measured using a GFTB 200 (Greisinger) or LM-9000 (Lutron), and sample temperatures were measured using a CT-321WP probe (CUSTOM). The pH of selected soil/sediment samples was measured using a LAQUAtwin pH-33B (Horiba, Kyoto, Japan). Excess liquid saturating soil/sediment was used to perform the measurements. The pH of pond water samples was measured similarly. For snow samples, pH was measured via a WQ330-PCD-S (Horiba) using pore water from snow cores or snow melt water. Dissolved oxygen concentrations were measured using liquid from samples using the WQ330-PCD-S, and conductivity was measured using the WQ330-PCD-S or a LAQUAtwin EC-33B (Horiba). In addition, total dissolved iron concentrations were measured using the WAK-Fe or WAK-Fe (D) kits (Kyoritsu Chemical-check lab, Yokohama, Japan), which rely on o-phenanthroline or bathophenanthroline, respectively, as the active reagent. Liquid from samples for iron concentration assays was passed through 0.45 μm filters before measurement.

*Environmental amplicon analysis*

Amplicon sequencing was performed by the Bioengineering Lab Co. (Sagamihara, Japan) using the PCR primers 341f (5'-CCT ACG GGN GGC WGC AG-3') and 805r (5'-GAC TAC HVG GGT ATC TAA TCC-3') with attached linking adapters. Subsequent sample processing, including sequencing on a MiSeq (2x300 bp; Illumina), relied on the same methods as the amplicon sequencing analysis described above for pure cultures, except the first round of PCR was performed using the KOD FX Neo polymerase (TOYOBO). Samples from 2022 and 2024 were processed on separate MiSeq runs. Sequencing generated 30,188-71,845 raw read pairs per sample.

Primers were trimmed from reads using the cutadapt trim-paired module including the '--p-discard-untrimmed' flag. Denoising was then performed using the dada2 denoise-paired module and was done separately for 2022 and 2024 samples given that these were sequenced independently. Sequence truncation lengths during denoising were determined based on quality score plots generated using the 'demux summarize' module, and lengths were set to 270 (forward) and 198 (reverse) for both denoising runs. After denoising, the resulting amplicon sequence variant (ASV) tables and ASV sequences for the two runs were combined using the feature-table merge and merge-seqs modules. To assign sequence taxonomy, a RESCRIPt-processed Silva r138 SSU database, clustered at 99% sequence identity [36, 107], was trimmed to the amplified 16S rRNA gene region using the feature-classifier extract-reads module, and the sequence classifier was then trained using the feature-classifier fit-classifier-naive-bayes module. Taxonomy assignment was then performed for the ASV sequences via the feature-classifier classify-sklearn module using the classifier trained above. A normalized

ASV table with bound taxonomy and sequence information was then produced using ampliwrangler version 0.2.0 (https://github.com/jmtsuji/ampliwrangler, DOI:10.5281/zenodo.15018024). After processing, samples contained 26,686-56,283 sequence counts. Detection limits of ASVs in a sample were calculated by dividing the minimum allowable sequence count (two reads) by the total post-denoising read counts for the sample.

Obtained ASVs were classified into potential *Crenothrix* clades by phylogenetic placement. All ASV sequences that were classified to the *Methylococcales* order with at least 0.3% relative abundance in any sample were analyzed alongside the collection of 16S rRNA gene reference and environmental sequences described above for phylogenetic analyses. The sequence set and ASVs were aligned using ssu-align version 0.1.1 [108], and the alignment was masked using the ssu-mask module. A phylogeny was built from the resulting multiple sequence alignment using IQ-TREE version 2.4.0 [100], with TIM+F+R4 being selected as the best-fit evolutionary rate model using ModelFinder [101] and with 1000 ultrafast bootstrap iterations [102]. Placement of the ASV sequences into monophyletic clades corresponding to Crenothrix-classified reference sequences (Fig. S2) was used to determine ASV classification.

To construct an ordination using the amplicon data, the diversity.beta_diversity function of scikit-bio version 0.6.2 [43] was used to generate a dissimilarity matrix, using Bray-Curtis distances, from the ASV table data in Python version 3.12.4. The was then generated as a PCoA using the stats.ordination.pcoa function of scikit-bio. Selected numerical metadata, including gas flux data and summed relative abundances of key taxa, were then normalized to lie between 0-1 using the preprocessing.MinMaxScaler function of scikit-learn version 1.5.1 [109] . These numerica metadata were used to calculate distance biplot values using scikit-bio's stats.ordination.pcoa_biplot function. For ease of viewing, the resulting biplot vectors were normalized so that the longest vector extended to the end of the plotted area of the ordination.

*CARD-FISH analysis*

The probes and formamide concentrations used in this study were Mγ669 (5'-GCT ACA CCT GAA ATT CCA CTC-3') [110] used at 20% formamide, Rhod662 (5'-GTC ACA AAT GCA GGT CCC AGG T-3')[111] used at 40%, and GEO3 A-C [112] used at 30% with helpers HGEO3-3mod (5'-GTT TAC GGC GTG GAC TAC C-3') and HGEO3-4 (5'-CAC TGC AGG GGT CAA TAC-3'), respectively. The GEO3 A-C probe was a mixture of A (5'-CCG CAA CAC CTA GTA CTC ATC-3'), B (5'-CCG CAA CAC CTA GTT CTC ATC-3'), and C (5'-CCG CAA CAC CTG GTT CTC ATC-3').
